# Structural basis of envelope and phase intrinsic coupling modes of the cerebral cortex

**DOI:** 10.1101/500009

**Authors:** Arnaud Messé, Karl J. Hollensteiner, Céline Delettre, Leigh-Anne Dell, Florian Pieper, Lena J. Nentwig, Edgar E. Galindo-Leon, Benoît Larrat, Sébastien Mériaux, Jean-François Mangin, Isabel Reillo, Camino de Juan Romero, Víctor Borrell, Gerhard Engler, Roberto Toro, Andreas K. Engel, Claus C. Hilgetag

**Author notes:** These authors contributed equally to this work (Andreas K. Engel, Claus C. Hilgetag).

## Abstract

Intrinsic coupling modes (ICMs) provide a framework for describing the interactions of ongoing brain activity at multiple spatial and temporal scales. Two families of ICMs can be distinguished: phase and envelope ICMs. The principles that shape these ICMs remain partly elusive, in particular their relation to the underlying brain structure. Here we explored structure-function relationships in the ferret brain between ICMs quantified from ongoing brain activity recorded with chronically implanted ECoG arrays and structural connectivity (SC) obtained from high-resolution diffusion MRI tractography. Large-scale computational models as well as simple topological ingredients of SC were used to explore the ability to predict both types of ICMs. Importantly, all investigations were conducted with ICM measures that are sensitive or insensitive to volume conduction effects. The results show that both types of ICMs are strongly related to SC, except when using ICM measures removing zero-lag synchronizations. Computational models are challenged to predict these ICM patterns consistently, and simple predictions from SC topological features can sometimes outperform them. Overall, the results demonstrate that patterns of cortical functional coupling as reflected in both phase and envelope ICMs bear a substantial relation to the underlying structural connectivity of the cerebral cortex.

## Introduction

Intrinsic coupling modes reflect the patterns of synchronization or functional connectivity (FC) between neuronal ensembles during spontaneous brain activity [1]. These coupling modes represent a widely used concept in modern neuroscience for probing the connectional organization of intact or damaged brains. ICMs have been associated with individual brain characteristics such as behavior and cognitive abilities [2–5], and aberrant ICM patterns have been linked to a variety of brain diseases [6–8]. Numerous studies in animals and humans have suggested that ICMs occur across a broad range of spatial and temporal scales, involving two distinct families of dynamical coupling mechanisms, namely phase ICMs (pICMs) and envelope ICMs (eICMs) [1]. Phase ICMs represent ongoing oscillations of band-limited dynamics, typically occurring at frequencies between about 1 to 150 Hz, which can be quantified by measures of phase coupling. By contrast, envelope ICMs correspond to coupled slow aperiodic fluctuations, typically on time scales of a few seconds to minutes, which can be uncovered by the correlation of signal power envelopes. However, the principles that shape both types of ICMs remain partly unresolved, notably their relation to the underlying brain structure [9,10]. Therefore, a careful investigation of the relation of ICMs to structural connectivity is required in order to advance our understanding of the network basis of cognition and behavior [10–15].

The physical basis for the emergence of functional connectivity patterns is the underlying brain anatomy, specifically, structural connectivity among brain regions [16]. In this context, diffusion MRI has transformed the field by its ability to estimate the whole brain architecture of structural connections in a non-invasive way [17]. In parallel, resting-state fMRI has become a popular approach for probing the functional organization of the brain at slow time scales, representing eICMs [18,19]. Pioneering studies have demonstrated a tight intricate association between the patterns of SC and resting-state fMRI-based FC [20–23]. Much less well understood are interactions at faster time scales, that is, pICMs, and their relations to the underlying anatomy. Nonetheless, a few studies have reported a positive association between pICMs and SC, using electrophysiological measurements [24,25]. However, a comprehensive description of the structure-function relationships including both types of ICMs and across frequency ranges is lacking.

Despite significant progress made in the analysis of functional neuroimaging data, such measurements suffer from diverse limitations. While fMRI has a good spatial resolution, its temporal resolution is rather limited, while the opposite is true for electrophysiological data. Furthermore, electrophysiological recordings are prone to present artefactual functional connectivity between close brain areas that is caused by the mixing of cortical signals at the sensors level due to volume conduction effects [26]. Because volume conduction is instantaneous (zero-lag), a number of measures quantifying only lagged couplings or strategies that explicitly remove zero-lag couplings have been designed to tackle this issue (‘lagged ICMs’) [27]. However their effects on the structure-function relationships remain poorly investigated (but see [28]).

Large-scale computational models have offered new mathematical tools to explicitly link SC to ICMs and to highlight generative mechanisms of functional coupling [29–31]. Most computational models presented in recent years have focused on describing functional interactions at slow time scales (eICMs) using resting-state fMRI [32]. A variety of models have been developed for this purpose that specifically consider the influence of a variety of parameters such as structural weights, delays and noise on neural dynamics [11,30,32]. Remarkably, substantially different computational models result in similar FC predictions [33, 34], in such a way that simple statistical models based on the topological characteristics of SC can be used with similar levels of prediction [35, 36]. Few studies have extended this framework by using electrophysiological measurements [37–39]. Recent investigations suggest that neuronal synchronization at faster time scales (pICMs) can also be predicted by computational simulations [28, 38]. Yet, the relative ability of computational models to predict both types of ICMs remains unanswered, especially when using lagged ICM measures. It is also unclear whether simple statistical models, based on SC topology, can equally predict phase as well as envelope ICMs. Overall, the mechanisms underlying the two types of ICMs are only partly resolved.

Against this background, we used data of ongoing activity of multiple cortical areas recorded from awake ferrets using chronically implanted micro ECoG arrays [40]. Such a setting offers recordings with high temporal and spatial resolution, for a substantial number of cortical areas. Moreover, due to the close proximity of the electrodes to the brain, volume conduction is strongly reduced, compared to non-invasive approaches such as M/EEG. Additionally, we obtained structural connectivity estimates from diffusion MRI tractography data for the regions underlying the ECoG array [41]. We found that both types of ICMs partially reflect the underlying SC across multiple frequency bands, and importantly, pICMs and eICMs show resemblance in their topographies, potentially pointing to a similar relation to the underlying SC. The use of lagged ICMs virtually abolishes the association with SC, since this removes zero-lag couplings which substantially contribute to local FC. Also, we found that computational models, independently of their complexity, can reproduce ICM patterns reasonably well. Furthermore, both types of ICMs can be predicted relying solely on the SC topology. Thus, the results demonstrate that patterns of cortical functional coupling as reflected in both phase and envelope ICMs are substantially constrained by the underlying structural connectivity of the cerebral cortex.

## Results

Intrinsic coupling modes were extracted from ECoG data of ongoing brain activity from awake ferrets [40]. Cortical activity was recorded in five adult female ferrets over multiple sessions using a 64 electrodes array distributed over half of the left cortical hemisphere. The animals’ brain states had been previously classified into slow-wave sleep, rapid-eye-movement sleep and awake periods using a data-driven approach [40]. The present study focused only on data from awake periods. For each animal and each awake period, pICMs and eICMs were computed in a frequency resolved manner (0.5–200 Hz) between all pairs of electrodes by quantifying coherence and power correlation, respectively. Then electrodes were assigned to cortical areas using the atlas of Bizley et al. [42] (Fig 1A), and the average ICM values between these areas were computed. Subsequently, ICM matrices were averaged within canonical frequency bands: 0.5-3 Hz (*δ*); 4-7 Hz (*θ*); 8-15 Hz (*α*); 16-30 Hz (*β*); 30-100 Hz (*γ*) (Fig 1B). ICM matrices were also averaged across awake periods within each session per animal (session-level), across sessions within each animal (animal-level), and also across animals (group-level). Results on the animal- and group-levels are presented in the main text, while results on the session-level are reported in supplementary figures. The corresponding structural connections of the areas covered by the ECoG array were assembled from diffusion MRI tractography data [41] (Fig 1C). Finally, in order to predict empirical patterns of ICMs, we employed computational biophysical models of various complexity as well as simple topological ingredients (Fig 1D). See Materials and Methods section for further details.

**Fig 1.**
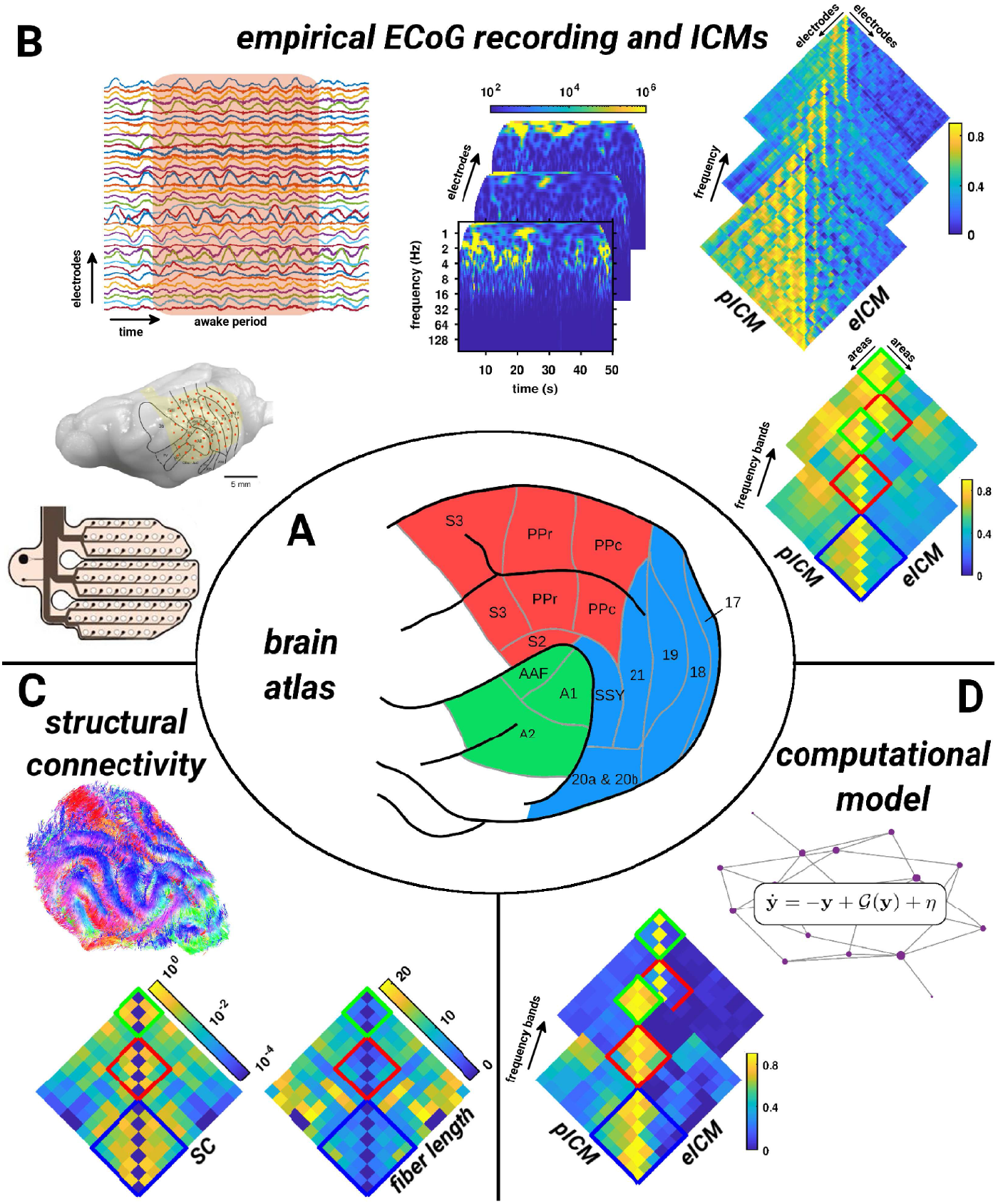
Multimodal processing pipeline. (A) Ferret cerebral cortex parcellated into 13 functionally and anatomically distinct regions according to [42]. Abbreviations: S2, S3: somatosensory cortex 2 and 3; PPr/c: rostral/caudal posterior parietal cortex; 17, 18, 19, 20, 21: occipital visual cortex areas; SSY: suprasylvian field; A1, A2: primary and secondary auditory cortex; AAF: anterior auditory field. (B) Empirical ECoG recording and ICMs. (From left to right) ECoG-array design and example of its placement over the left posterior ferret brain cortex. The ECoG consists of 64 platinum electrodes distributed across three polymide fingers to enable the array to adapt to the curved surface of the cortex. Awake periods were extracted from the resulting brain activity, then time-frequency representations were computed using wavelet transforms for each electrode. Phase and envelope ICMs were computed over multiple frequencies and between all pairs of electrodes. Subsequently, ICMs were mapped to the ferret brain parcellation and averaged within canonical frequency bands. (C) Patterns of structural connectivity extracted from diffusion MRI tractography and the associated fiber lengths. (D) Large-scale computational brain models and an illustrative example of simulated phase and envelope ICMs across frequency bands.

### Structure-function relationships

The similarity between ICM patterns and the underlying structural connectivity was quantified, for each animal and the group average, and for each frequency band, by means of Spearman correlations between SC and phase and envelope ICM values. Phase and envelope ICMs were both significantly positively correlated to SC in a similar way for all frequency bands, except for one animal for both ICM measures across all frequency bands (Fig 2A). The correlations of phase and envelope ICMs with SC were not significantly different across all frequency bands for all animals and in the group average. Next, we evaluated the influence of the physical distance between areas in the structure-function relationships. As expected, both SC and ICMs were significantly negatively correlated with distance. Similar for both, connectivity values decrease with increasing inter-areal distance, except for one animal for both pICM and eICM in the *δ* band (Fig 2A). Of note, SC is more strongly influenced by distance than are ICMs. Consequently, the correlation between SC and ICMs are largely reduced when controlling for distance. Two animals had no significant partial correlation of pICM with SC for all frequency bands except the *γ* band, where pICM appeared significantly negatively partially correlated with SC. One animal (resp. the group average) showed no significant partial correlation of pICM with SC in the δ (resp. γ) band. Regarding eICMs, one animal had no significant partial correlation with SC for all frequency bands except the *β* and γ bands, where eICMs appeared significantly negatively partially correlated with SC. One animal and the group average showed no significant partial correlation with SC for all frequency bands (Fig 2A).

**Fig 2.**
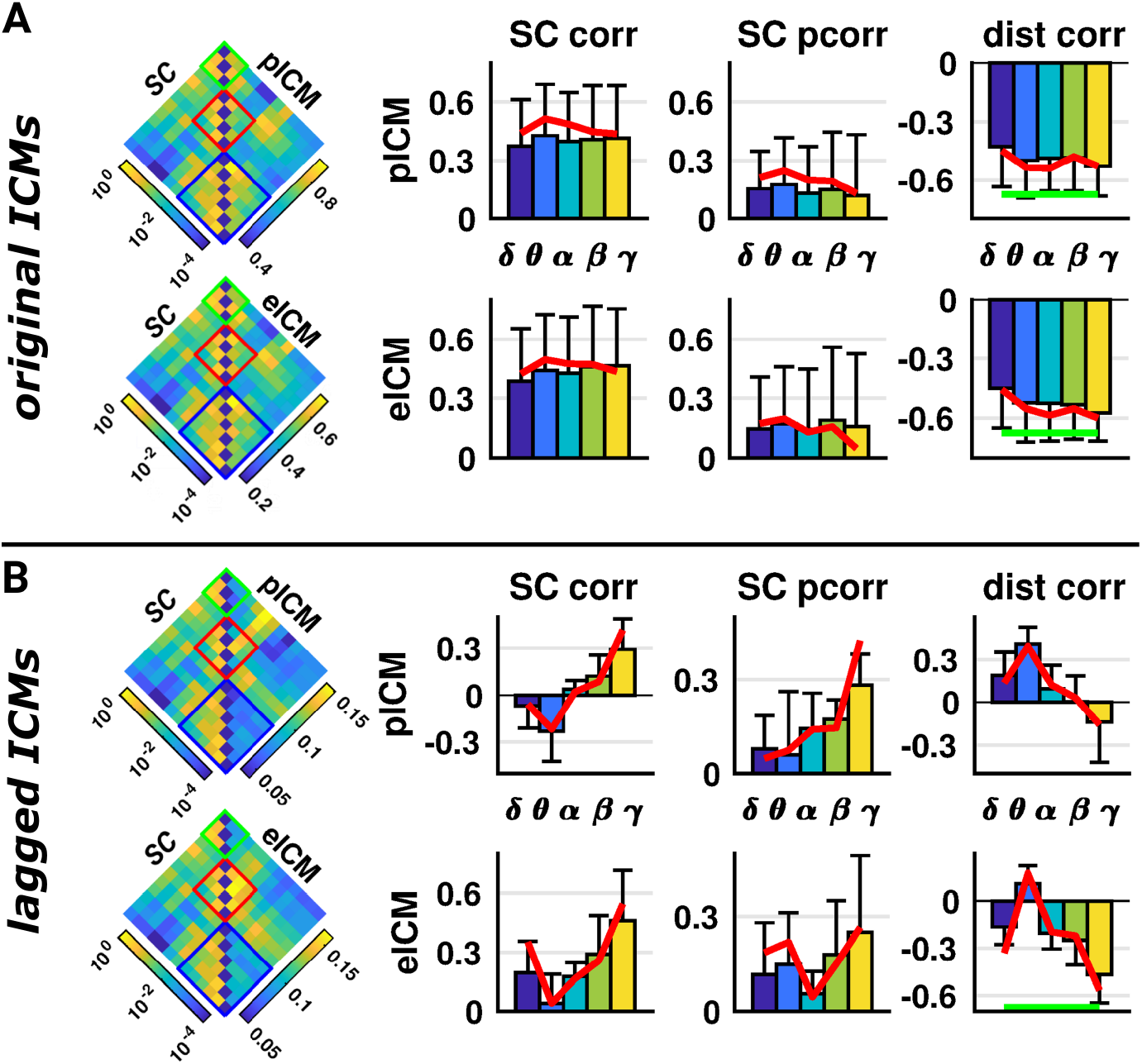
Structure-function relationships. (A) Correlation between structural connectivity and phase and envelope ICMs across frequency bands and animals. (Left) Patterns of SC (left triangular part) and group level phase or envelope ICMs (right triangular part) for the α band. (Right) For each ICM measure and each frequency band, we represented the individual correlations (bar chart representing means and associated standard deviations), as well as the correlation for the group average (red curve), between SC and ICMs (SC corr), between SC and ICMs when controlling for the distance (SC pcorr), and between distance and ICMs (dist corr). The green line represents the correlation between distance and SC. (B) Same as (A) when using lagged ICM measures.

In addition, in order to deal with the potential confounds resulting from volume conduction, we also computed the correlation between SC and lagged ICM measures (Fig 2B). Envelope ICMs were significantly positively correlated to SC for three animals and the group average for all frequency bands except the *θ* and *α* bands, two animals had non significant correlations for all frequency bands (Fig 2B). Phase ICMs were found to be only significantly positively correlated to SC in the *γ* band for three animals and the average, and negatively correlated to SC for the *θ* band in two animals (Fig 2B). The correlation of pICMs with SC was significantly lower compared to eICMs for low frequency bands in few animals. We observed an increase in the correlation of both types of ICMs with SC with increasing frequency. Phase (resp. envelope) ICMs were significantly positively (resp. negatively) correlated with distance for the low (reps. high) frequency bands (Fig 2B). Consequently, the correlation between SC and pICMs remained significant only for the *γ* band, while the correlation between SC and eICMs remained significant only for low and high frequency bands in few animals when controlling for distance (Fig 2B). The partial correlations of phase and envelope ICMs with SC were not significantly different. For a complete description of the significant statistical tests see Fig S1 and for the results at a finer level see Figs S2 and S3.

### Phase and envelope ICMs similarity, and interfrequency, interindividual consistency

The similarity between pICMs and eICMs was quantified, for each frequency band, each animal and the group average, by means of Spearman correlation. Phase and envelope ICMs were significantly positively correlated with each other in all frequency bands for all animals (Fig 3A). Consistency of ICM measures across frequency bands was quantified, for each pair of frequency bands, each animal and the group average, by means of Spearman correlation. We observed a significant high interfrequency consistency for each ICM measure which is consistent across animals (Fig 3A). Of note, the similarity was higher between ICM measures within frequency bands than between frequency bands within ICM measures. We also assessed the consistency of both ICM measures across animals, for each pair of animals, and each frequency band, by means of Spearman correlation. Interindividual consistency was significant across all animals and for both ICM measures (Fig 3A). We observed a slight increasing interindividual consistency with increasing frequency.

**Fig 3.**
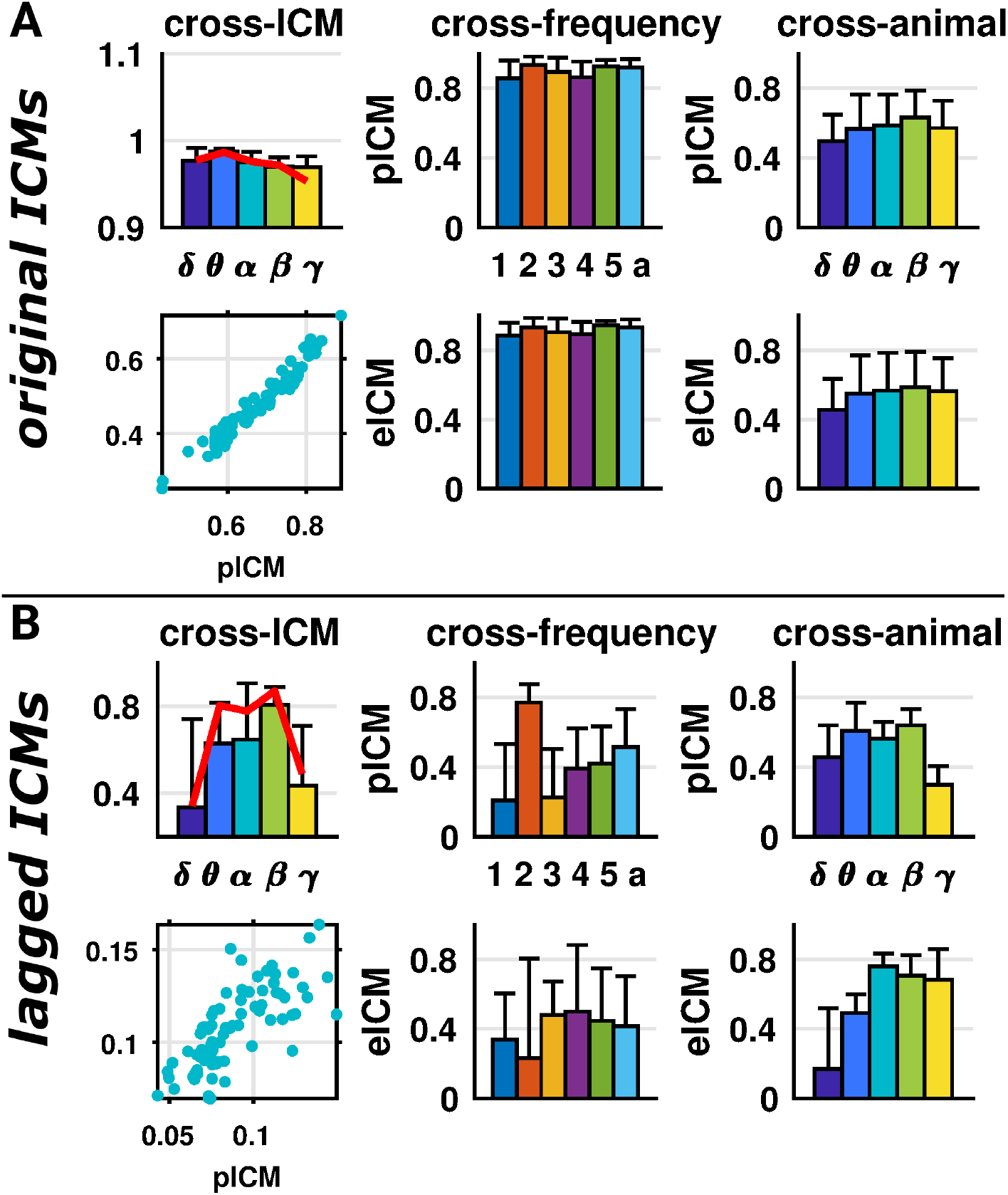
ICM similarity and consistency. (A) Similarity of phase and envelope ICMs, and consistency across frequency bands and animals. (Left) Correlation between phase and envelope ICMs across frequency bands and animals. (Top) For each frequency band, we represented the individual correlations (bar chart representing means and associated standard deviations), as well as the correlation for the group average (red curve). (Bottom) Example scatter plot of phase versus envelope ICM values for the group average in the α band. (Middle) Correlation of phase (top) and envelope (bottom) ICM patterns between frequency bands. For each ICM measure, each animal (1 to 5) and the group average (a), we represented all pairwise correlations between frequency bands (bar chart representing means and associated standard deviations). (Right) Correlation of phase (top) and envelope (bottom) ICM patterns between animals. For each ICM measure and each frequency band, we represented all pairwise correlations between animals (bar chart representing means and associated standard deviations). (B) Same as (A) when using lagged ICM measures.

When lagged measures phase and envelope ICMs were used, consistent results were observed (Fig 3B). Phase and envelope ICMs were significantly positively correlated with each other for most frequency bands and animals, albeit to a much lower level compared to the original ICM measures, especially for the *δ* and γ bands (Fig 3B). Interfrequency consistency was significant across most animals and pairs of frequency bands, albeit at a much lower level compared to the original ICM measures (Fig 3B). Interindividual consistency was significant across all pairs of animals, all frequency bands, and for both ICM measures, except in the *δ* band for eICMs (Fig 3B). Of note, the consistency of phase (resp. envelope) ICMs in the *γ* (resp. *δ*) band was largely reduced compared to the original ICM measures. For a complete description of the significant statistical tests see Fig S4 and for the results at a finer level see Figs S5 and S6.

### Computational model predictions

Next, we employed computational models of various complexity for predicting ICM measures: the spiking attractor network (SAN) model, a biologically realistic model of a large network of spiking neurons; the Wilson-Cowan (WC) model, a popular neural-mass model of coupled excitatory and inhibitory populations; and the spatial autoregressive (SAR) model, a statistical model capturing the stationary behavior of a diffuse process on networks. We quantitatively fit the models using different optimization strategies to explore the models’ ability to predict ICM measures (see Materials and Methods section for details). Model parameters were chosen to minimize an objective function, defined as the distance between simulated and empirical ICMs averaged over frequency bands. We either minimized the objective function averaged across all ICM measures combining both original and lagged measures (one parameter set; *all*), over only original (*original*) or only lagged (*lagged*) ICM measures (two parameter sets), or for each ICM measure individually (four parameter sets; *individual*).

Parameter space exploration highlights the sensitivity of the objective function to model parameter values, the global coupling strength and mean delay (Fig 4). None of the models possessed parameter settings which can minimize the distance of the simulations to all ICM measures simultaneously. Optimizing all measures simultaneously or only the original ICM measures led to a similar set of parameter values, except for the SAR model. The optimization of the lagged ICM measures generally led to parameter values far from the ones for the original ICM measures.

**Fig 4.**
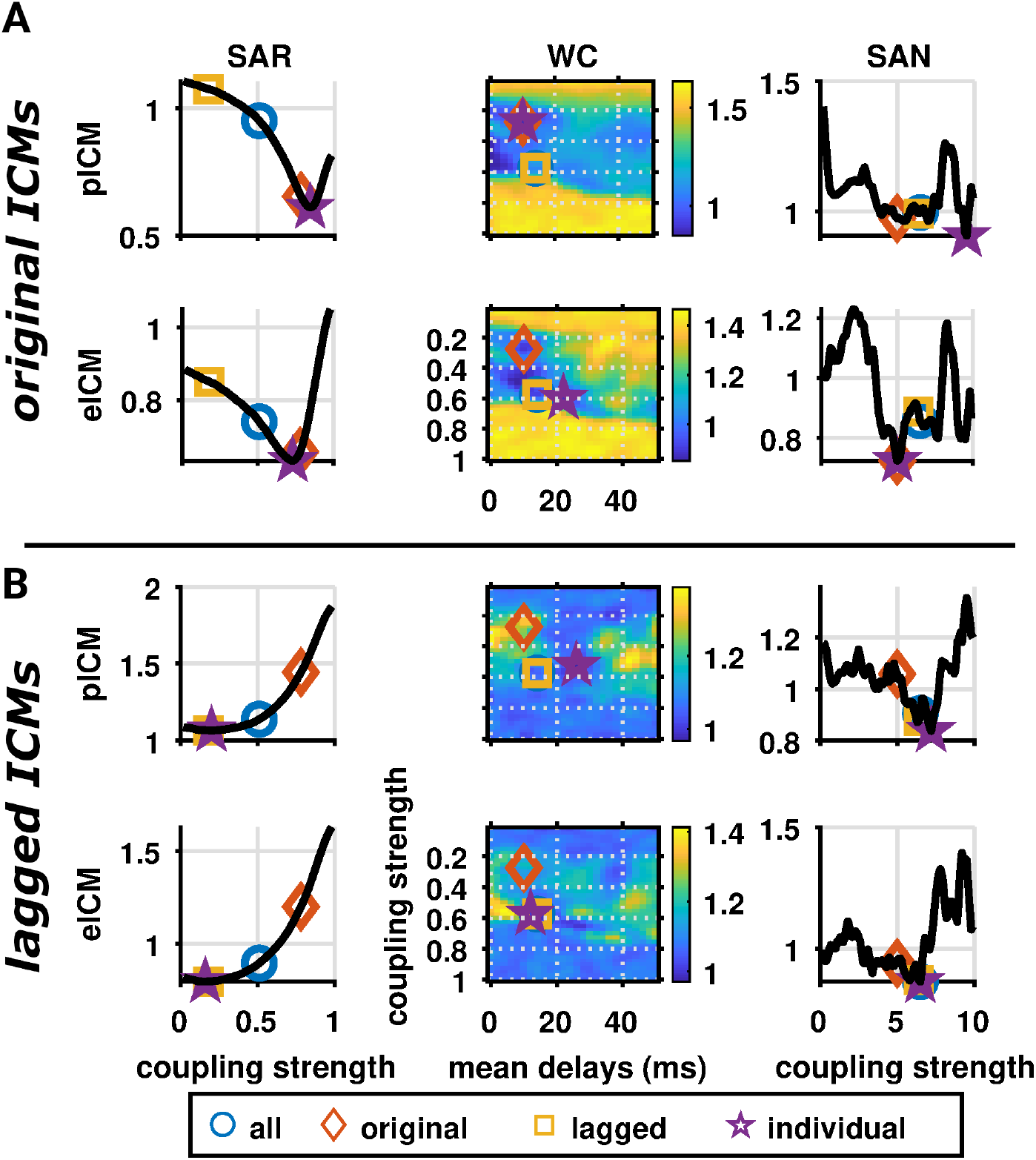
Parameter space exploration of the computational models. (A) Objective function values between simulated and empirical phase and envelope ICMs as a function of the models’ parameters when using original ICM measures. Columns represent the different models (left: SAR; middle: WC; and right: SAN). Parameter values were chosen to either minimized the objective function averaged across all ICM measures combining both original and lagged measures (blue circle, *all*), over only original (red diamond, *original*) or only lagged (yellow square, *lagged*) ICM measures, or for each ICM measure individually (purple star, *individual*). (B) Same as (A) when using lagged ICM measures.

In general, computational models predicted phase and envelope ICMs in a similar way (Fig 5). Most of the correlation between simulated and empirical ICMs were significantly positive, except when using lagged ICM measures, where few correlations were significant for pICM and correlations were mostly significant for high frequency bands for eICM (Fig S7). Simulated ICMs were not significantly more correlated to the empirical data compared to SC alone (Fig S8). Even when optimizing specifically lagged ICM measures (*lagged* and *individual*), no model seemed able to predict these ICM patterns accurately. We observed only positive correlations for high frequency bands (*β* and *γ*). Overall, these results highlight the challenge of properly fitting biophysical models to rich empirical datasets.

**Fig 5.**
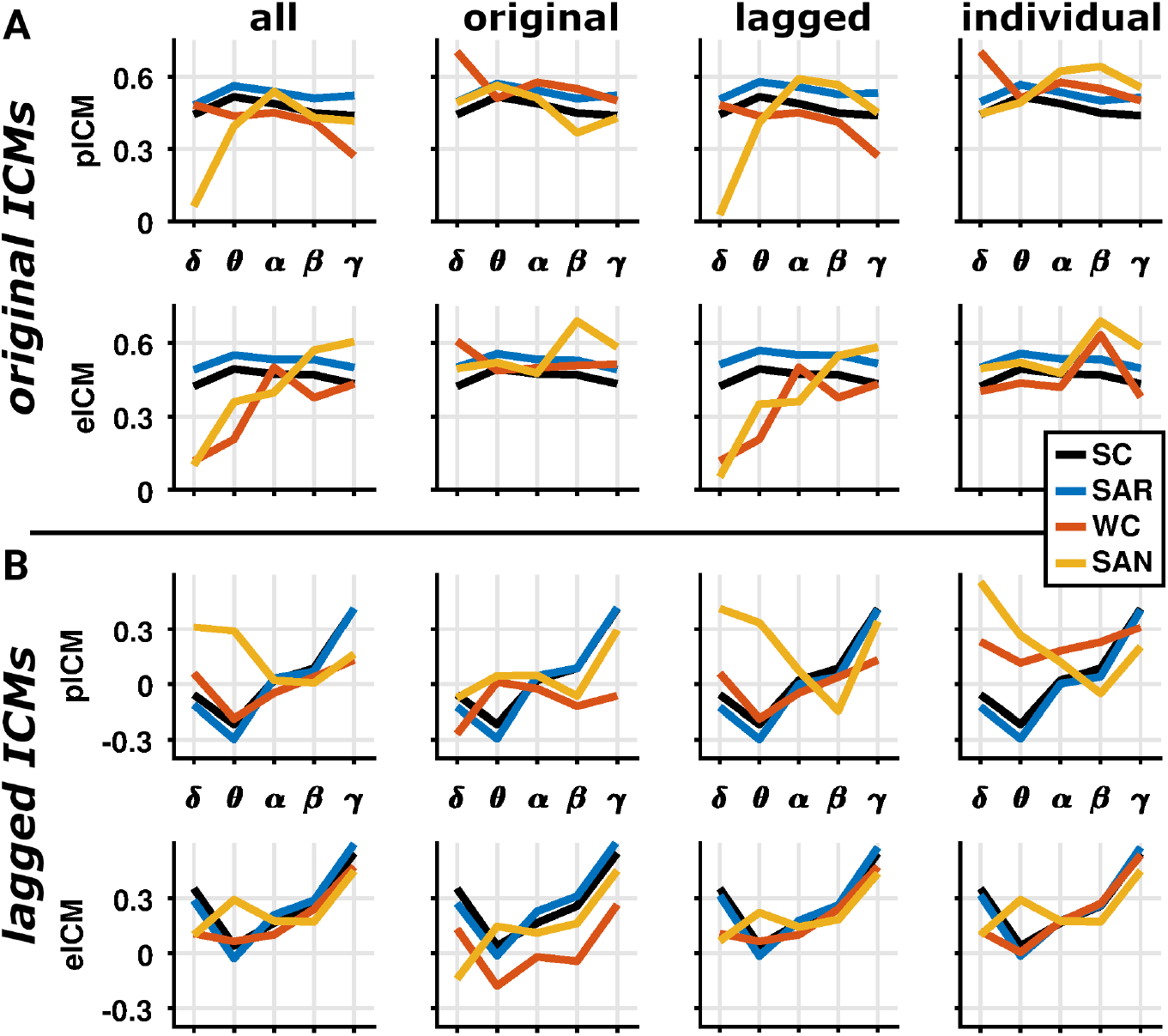
Predictive power of the computational models. (A) Correlation between simulated and empirical phase and envelope ICMs across frequency bands and models when using original ICM measures. Columns represent the different optimization strategies. We either minimized the objective function averaged across all ICM measures combining both original and lagged measures (first column, *all*), over only original (second column, *original*) or only lagged (third column, *lagged*) ICM measures, or for each ICM measure individually (last column, *individual*). (B) Same as (A) when using lagged ICM measures.

### Topological prediction

To better understand the emergence of ICMs from SC and to circumvent the challenge of properly calibrating computational models, we looked at the relationships of the ICMs with the underlying SC topological architecture. The most prominent topological feature that facilitates communication in networks is captured by structural walks, that is the ordered sequence of links connecting areas. Using spectral theory properties, we computed the total weight of all the walks of a given length linking all pairs of areas (see Materials and Methods section for details). Subsequently, the similarity between ICMs and structural walks was quantified, for each animal and the group average, and for each frequency band, by means of Spearman correlation between SC walk weights and phase and envelope ICM values. Phase and envelope ICM measures were both similarly positively correlated with the weights of the walks, but the correlation decreased with increasing walk length and vanished for walks of length greater than 3 (Fig 6 and S9). Little variations were observed across frequency bands. When looking at lagged ICM measures, the pattern of associations is almost inverted. Lagged ICMs were more strongly correlated with longer walks, especially for low frequency bands. Interestingly, using a weighted linear combination of the walks’ weights allowed us to predict original ICM patterns well, but somewhat less for the lagged ICM measures (Fig 6). While the prediction of the original ICM measures reached values close to 0.7, the prediction of lagged ICM measures rarely exceeded 0.6 (for comparison, the models’ prediction was at best of 0.3). For a complete description of the significant statistical tests see Fig S10.

**Fig 6.**
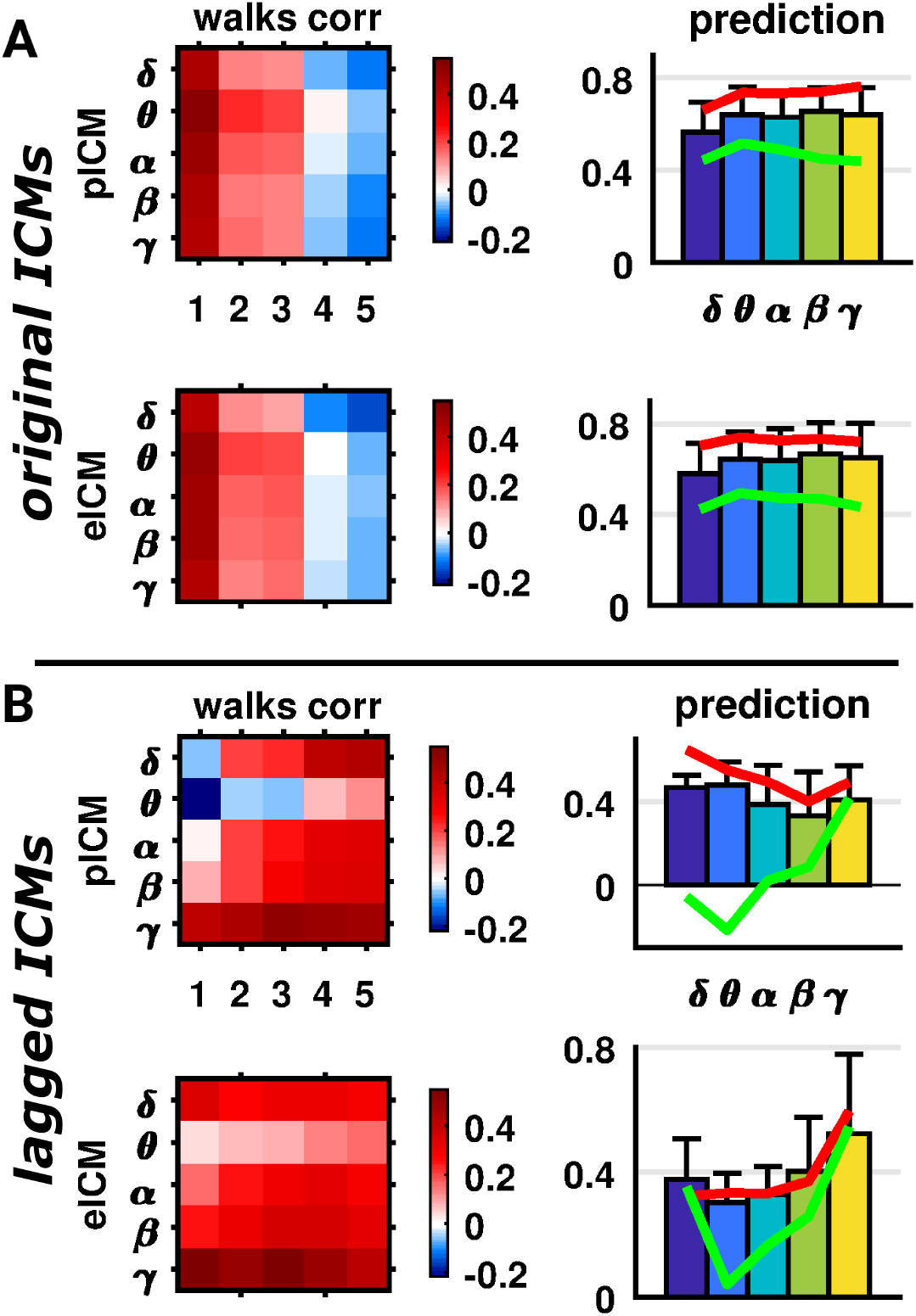
Predictive power of the structural walks. (A) Correlation between structural walks and empirical phase and envelope ICMs across frequency bands. (Left) For each ICM measure, each walk length and each frequency band, we represented the correlation for the group average. (Right) For each ICM measure and each frequency band, we represented the individual correlations (bar chart representing means and associated standard deviations), as well as the correlation for the group average (red curve), between the linear combination of the walks and ICMs. We additionally plotted the correlation between SC and ICMs for the group average (green curve) as reference. (B) Same as (A) when using lagged ICM measures.

## Discussion

### Overview

In the present study, we investigated the relationship between structural connectivity and ICMs of ongoing dynamics in the ferret brain. We took advantage of the high temporal resolution offered by ECoG measurements to explore these relationships across multiple frequency bands and different animals. We also investigated the similarity between phase and envelope ICMs across frequency bands, as well as their interfrequency and interindividual consistency. Additionally, we explored and compared the performance of computational models of various complexity and some topological ingredients in predicting simultaneously envelope and phase ICMs. All investigations were conducted with ICM measures sensitive or insensitive to volume conduction effects. Generally, we found that SC is significantly related to both types of ICMs in a similar way. When using original ICM measures both pICMs and eICMs are highly similar and consistent across frequency bands and animals. Using lagged ICM measures virtually abolished the association with SC, and reduced their similarity and consistency. Computational models are challenged to predict these ICM patterns consistently and simultaneously. The models are generally good at predicting original ICM measures but fail for the lagged ones. Interestingly, the emergence of ICM patterns seems largely driven by the underlying SC topological architecture, where a linear combination of structural walks outperformed the prediction compared to the employed computational models. Overall, our results suggest that ICMs during ongoing activity in awake ferrets reflect, to a significant degree, the topological organization of anatomical cortical networks, but are likely also influenced by other factors.

### Structure-function relationships

In previous studies, envelope ICMs and their association with structural connectivity were mainly investigated using resting-state fMRI data, where consistent positive correlations between SC and eICMs were reported [20–23,33,43], reviewed in [44]. Some recent studies have extended and confirmed these observations based on the refined temporal resolution offered by electrophysiological measurements. The relationship between pICMs and SC has been explored using both EEG [24, 28, 45] and MEG [25, 46] data, while eICMs were investigated using MEG recordings [39, 47]. Similar to the resting-state literature, a positive relationship between both types of ICMs with SC has been reported consistently, but its potential frequency dependence has not been thoroughly investigated [24, 47]. Here, we confirm these observations by showing that both types of ICMs are positively correlated to SC and this across the frequency ranges. As a matter of fact, both measures appeared to be also correlated to each other independently of the frequency. Additionally, we observed a relatively high consistency of the original ICM measures both across frequencies and animals, similar to observations in MEG recordings [48].

One important confound in the analysis of functional connectivity using electrophysiological measurements is the potential presence of volume conduction. A number of measures and procedures have been designed to mitigate such effects. Most of them specifically remove zero-lag synchronization or coactivations [49]. But, reliable estimation of ICMs with electrophysiological recordings remains challenging due to signal mixing [50, 51]. We observed that the similarity between phase and envelope ICMs, and their consistency across frequencies are largely reduced when using lagged measures, as previously reported [48, 52]. The correlation between both types of ICMs with SC appears also largely reduced and virtually abolished, except for high frequency, compared to the original measures. Such results showed us that the classical ICM measures are strongly driven by near-zero coactivations, which are in part resulting from volume conduction, but in part also due to physiological synchrony [1, 2, 7], which occurs most prominently across short distances and depends on the underlying network structure [53].

### ICMs models

Computational brain models give us a parsimonious approach to explore the generative mechanisms of ICMs [31, 32, 54]. Most modeling studies have investigated the emergence of eICMs using resting-state fMRI [11,33,34,55] or MEG data [37–39], while few studies have explored the generative processes of pICMs using MEG [38] or EEG [28] recordings. Using computational models of various complexity, we showed their potential for predicting both types of ICM. None of the models investigated here appears to be able to predict simultaneously original and lagged ICM measures. As a matter of fact, none can predict lagged measures, even when fitting the models explicitly for these measures. One of the most intriguing results is the fact that none of the models seems to predict ICM patterns much better than SC alone. Looking at the topological architecture of SC and especially the structural walks, we showed that original ICMs are mostly explained by short walks while the opposite is true for the lagged measures. As such, and as previously observed in resting-state fMRI studies, patterns of functional interactions appear substantially constrained by the underlying topological scaffold of the brain [35, 56].

### Biological significance

On the one hand, the removal of zero-lag interactions avoids artefacts that is known to arise from signal mixing at sensor level [12, 57, 58]. On the other hand, such a treatment may also remove actual biological signals. Indeed, the present results confirm that zero-lag interactions are an essential component of cortical coupling modes [1,2,7,13]. Adjacent regions of the ferret cortex are strongly connected by short fibres [41]. This kind of connectivity supports very fast, zero-lag interactions between neighbouring regions [59]. The importance of these interactions for intrinsic cortical coupling is demonstrated by the observation that (original) eICMs as well as pICMs strongly depend on the physical embedding in the cortex and decay with distance (Fig 2A). Moreover, direct structural connections (walks of length one) are most strongly correlated with original eICMs and pICMs, while the correlation becomes gradually weaker for longer walks with longer delays (Fig 4A). Conversely, the dominant role of direct, short-distance interactions is abolished for lagged ICMs (Figs 2B, 4B). Instead, such ICMs are based more equally on walks of all lengths (Fig 4B), that is, mostly on indirect connectivity. Moreover, as structural cortical connectivity is dominated by short connections, the predictive power of computational models is much reduced for lagged versus original ICM measures. These findings once again underline the role of direct and indirect structural connectivity in cortical communication.

### Caveats

The present results are subject to several methodological considerations. First, the spatial coverage of the ECoG-array was limited to roughly one half of a cerebral hemisphere that limits us for a potential generalization and prevents us to study, for example, interhemispheric structure-function relations which have been studied in other preparations before [60] but shown to be challenging [33]. Additionally, we solely focused on the awake resting-state periods in order to make concrete comparisons with the existing literature mostly based on that brain state. Little is known about the eventual modulation of the structure-function relationships across diverse contexts, including, for example, sleep stages or when animals interact with the environment, deserving further investigations [52, 61]. Regarding computational modelling, we made use of standard models together with their default settings (except for the coupling strength and the average delay). For further refinement, one should consider to optimize a larger number of parameters or even to incorporate knowledge from external modalities including, for example, receptor maps [62] or anatomical laminar details [63].

## Materials and methods

### ECoG data

Intrinsic coupling modes were extracted from ECoG recordings of ongoing brain activity in awake ferrets (*Mustela putorius furo).* Data were collected in five adult female ferrets [40]. All experiments were approved by the Hamburg state authority for animal welfare and were performed in accordance with the guidelines of the German Animal Protection Law.

To obtain the recordings from an extended set of cortical areas in freely moving animals, an ECoG array had been implanted that was co-developed with the University of Freiburg (IMTEK, Freiburg) [64]. The array covered a large portion of the posterior, parietal, and temporal surface of the left ferret brain hemisphere.

Sixty-four platinum electrodes (*ø* 250 μm) were arranged equidistantly (1.5 mm) in a hexagonal manner. See Fig 1 for a schematic diagram of the ECoG layout. Ferrets had been accustomed to a recording box (45×20×52 cm), where they could move around freely while neural activity was recorded. For each animal, at least 4 separate recording sessions with a duration of at least 2 hours had been obtained. ECoG signals had been recorded using a 64 channel AlphaLab SnRTM recording system (Alpha Omega Engineering, Israel), filtered (0.1–357 Hz bandpass), and digitized (1.4 kHz sampling rate). A total of 23 recording sessions were obtained and analysed. The position of ECoG grids was then projected onto a scaled illustration of the atlas of Bizley et al. [42]. Data from each electrode were then allocated to the cortical area directly underlying the corresponding ECoG contact.

Noise epochs were detected using a threshold of 10 standard deviations. Data were rejected in a window of ± 10 s from all time points that exceeded this threshold. Then, notch filters were applied to remove line noise and its harmonics (50, 100, and 150 Hz), and the data were downsampled to 700 Hz. All data were visually inspected before further analysis to exclude electrical artefacts. In a previous study, brain activity was objectively classified into slow-wave sleep, rapid-eye-movement sleep and awake periods, using a data-driven approach [40]. The present study focused only on data from awake periods, on average, 34 periods of 78 s per session across animals (periods shorter than 10 s were discarded to increase statistical power). For each animal and each period, a time-frequency estimate X¿(t, *f*) of signal x¿(t) at electrode *i* was computed by convolution with a series of Morlet’s wavelets. Carrier frequencies were spaced logarithmically from 0.5 to 200 Hz. The width of the wavelets was set to 7 cycles. Spectral estimates were computed at a rate of 5 Hz. Then, phase and envelope ICMs were computed between all pairs of electrodes using the measures of coherence and power correlation, respectively, see Intrinsic coupling modes section.

Empirical ICM measures were first computed at the electrode level. To account for inter-animal differences in the position of the ECoG-array and thus make cross-animal comparisons possible, ICM matrices were calculated at the area-level using the atlas of Bizley et al. [42]. Each electrode was assigned to a cortical area by a ‘winner-takes-all’ method using its maximum overlap, and the average ICM values between these areas were computed. In total, 13 areas were covered by the ECoG-array for all animals (Fig 1). Subsequently, ICM matrices were averaged within canonical frequency bands: 0.5-3 Hz (*δ*); 4-7 Hz (*θ*); 8-15 Hz (*α*); 16-30 Hz (*β*); 30-100 Hz (γ) for condensed representations. Finally, ICM matrices were averaged across awake periods within each session per animal (session-level), then across sessions within each animal (animal-level), and also across animals (group-level). Results on the animal- and group-levels are presented in the main text, while results on the session-level are reported in supplementary figures.

All data analysis was performed using custom scripts for Matlab (Mathworks Inc) and the Fieldtrip software [65].

### Intrinsic coupling modes

Phase and envelope ICMs were estimated pairwise using the measures of coherence and power envelope correlation, respectively. Coherence represents the analogue of the classic correlation for frequency resolved data. Coherence is based on the normalized cross spectral density (or coherency), it is a measure of the linear relationship between oscillatory signals, where a high value corresponds to signals with similar amplitudes and aligned phases. The coherency between signals at location *i* and *j* and for frequency *f*, is given by

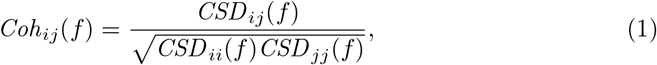

where 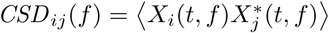 is the cross spectral density, with () the expected operator in time and * the complex conjugate. The measure of coherence is defined as the absolute value of the coherency, |*Coh*|.

Power correlation is a measure of the similarity between the power envelopes of the recorded signals [12,66,67]. Power envelope is given by the squared absolute values of the complex spectral estimates. Furthermore, a logarithmic transform was applied to render the statistics more normal. Then, the Pearson correlation between the resulting power envelopes from two different locations *i* and *j* was computed

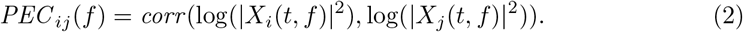

Alternatively, we also explored measures robust to volume conduction effects. Phase and envelope ICMs were estimated pairwise using the imaginary coherence and orthogonal power envelope correlation, respectively. Imaginary coherence, the imaginary part of coherency, 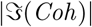, eliminates the contribution of volume-conducted signals by exclusively considering phase-lagged signal components [57]. Orthogonal power envelope correlation, similar to imaginary coherence, removes instantaneous covariations by orthogonalizing signals before computing their power envelopes [12,58]. The time-frequency estimate *X_i_*(*t, f*) orthogonalized with respect to signal *X_j_* (*t, f*) is given by

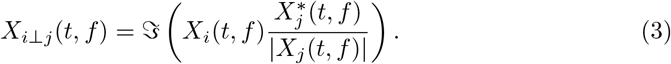

The orthogonalization being not symmetric, 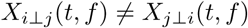, power envelope correlation was computed in both directions and subsequently averaged.

### Structural connectivity

Ferret brain structural connectivity data were based on diffusion MRI tractography covering the 13 areas available from the ECoG recordings [41]. High resolution MRI were acquired *ex vivo* from a *postmortem* 2 month old ferret using a small animal 7 Tesla Bruker MRI scanner (Neurospin, Saclay, France). The ferret was euthanized by an overdose of pentobarbital and perfused transcardially with 0.9% saline solution and post-fixed with phosphate-buffered 4% paraformaldehyde (PFA). After extraction, the brain was stored at 4°C in a 4% PFA solution until the MRI acquisition. All procedures were approved by the Institutional Animal Care and Use Committee of the Universidad Miguel Hernandez and the Consejo Superior de Investigaciones Científicas, Alicante, Spain.

High resolution T2-weighted MRI data was acquired using a multislice multiecho sequence with 18 echo times and 0.12 mm isotropic voxels. Diffusion MRI data were acquired using a multislice 2-D spin-echo segmented EPI sequence (4 segments) with the following parameters: TR = 40 s; TE = 32 ms; matrix size = 160×120×80; 0.24 mm isotropic voxels; 200 diffusion-weighted directions with b = 4000 s/mm^2^; and 10 b0 at the beginning of the sequence, diffusion gradient duration = 5 ms and diffusion gradient separation = 17 ms. The total acquisition time of the diffusion MRI sequences was about 37 hr.

Diffusion MRI were first visually inspected to exclude volume with artefacts. Then, the following preprocessing steps were carried out: local principal component analysis denoising [68], Gibbs ringing correction [69], FSL-based eddy current correction [70,71] and B1 field inhomogeneity correction [72]. A brain mask was manually segmented from the high-resolution T2 volume. Spatial normalization using a linear transformation between the high-resolution T2 volume and diffusion MRI data was computed using FLIRT tools [73], and the brain mask was registered to the diffusion space. Tractography was performed based on the fiber orientation distribution estimated with a multishell multitissue constrained spherical deconvolution (msmt CSD) [74]. Spherical harmonic order was set to 8. The response functions were computed using the ‘dhollander’ algorithm which provides an unsupervised estimation of tissue-specific response functions. The msmt CSD was performed using a WM/CSF compartment model [75]. The streamline tractographies were then produced following a probabilistic algorithm (iFOD2) [76]. One million streamlines were tracked over the full brain with the parameters recommended by MRtrix3: stepsize 0.12 mm, angle 45° per voxel, minimal streamline length 1.2 mm, maximal length 2.4 cm. Streamline seeds were produced at random locations within the brain mask until the defined number of streamlines was reached. To prevent streamlines from going across sulci, the brain mask was used as a stopping criterion.

A structural connectivity matrix was extracted from the tractography output using the number of streamlines connecting pairs of regions of a parcellation based on the atlas of the posterior cortex by Bizley et al. [42]. The parcellation scheme was manually drawn on the left hemisphere in the diffusion MRI space using the online tool BrainBox (http://brainbox.pasteur.fr/). The structural connectivity matrix was inherently symmetric as diffusion MRI tractography does not provide any information about directionality. A matrix reporting the averaged fiber lengths between regions was also computed. The structural connectivity network is represented by the weighted matrix *A,* where the entries aj are the weights of the connections, i.e., the number of streamlines, between pairs of areas i and j.

All data analysis was performed using custom scripts for Python (http://www.python.org/) and the MRtrix3 software (http://www.mrtrix.org/).

### Computational models

We employed computational models of various complexity: the spiking attractor network model, a biologically realistic model of a large network of spiking neurons [77], the Wilson-Cowan model, a popular neural-mass model of coupled excitatory and inhibitory populations [78], and the spatial autoregressive model, a statistical model capturing the stationary behavior of a diffuse process on networks [36]. All models incorporated a parameter that represents the coupling strength between regions. This parameter was optimized separately for each model, see Statistics section. The structural connectivity matrix was normalized before simulations such that the matrix’s rows sum to one [33, 79, 80].

### Spiking attractor network model

The SAN model is a detailed computational model based on spiking neurons and realistic synapses. A description for the microscopic level, that is within areas, is achieved by using a biologically realistic model of interconnected populations of excitatory and inhibitory neurons. The postsynaptic activity is dependent on the incoming synaptic input currents from other neurons through AMPA, NMDA and GABA receptors, as well as from external background input modelled by Poisson spike trains. Excitatory populations are interconnected at large-scale (inter-areal) *via* the structural connectivity scaled by a global coupling strength factor. For a detailed description of the model refer to [77], all parameter values (except the coupling strength) were retained from the original study.

We explored coupling parameter values from 0.1 to 10 by steps of 0.1. For each coupling strength value, the SAN model was simulated at a sampling frequency of 10 kHz for 5 minutes. The resulting data were then downsampled to 1 kHz. Simulated ICMs from the neuronal activity were extracted in a similar way as for the empirical data from a time-frequency decomposition of the simulated signals.

### Wilson-Cowan model

The WC model represents a network of neural masses describing the activity of an ensemble of excitatory (*E*) and inhibitory (*I*) neuronal populations [30,78]. The dynamics is governed by the following equations:

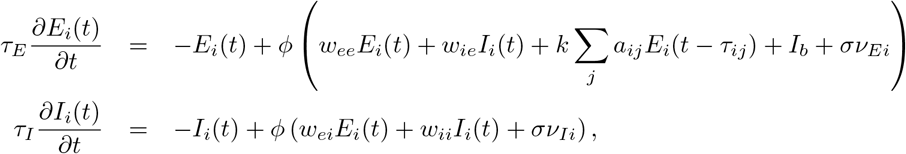

where *E_i_* (resp. *I_i_*) represents the average firing rate of the excitatory (resp. inhibitory) population *i*, and *τ_E_* (resp. *τ_I_*) the excitatory (resp. inhibitory) time constant. *w_ab_* is the local connectivity strength between populations *a* and *b,* and *k* represents the global coupling strength. *A* = {*a_ij_*} represents the structural connectivity matrix. *I_b_* is a constant spontaneous background input. *τ_ij_* is the propagation delay between regions *i* and *j*, based on the average fiber tract length between regions scaled by the axonal velocity, *v*, i.e., *τ_ij_* = *L_ij_*/*v. ν* is a random fluctuating input accounting for sources of biophysical variability, scaled by *σ*. The transfer function *ϕ* accounts for the saturation of firing rates in neuronal populations and is modeled by a sigmoid: *ϕ*(*x*) = [1 + *e*^-(*x-a*)/*b*^]^-1^. All parameter values (except the coupling strength and the delay) were set as in [38].

We explored coupling strength values from 0.025 to 1 by steps of 0.025, and mean delay values from 0 to 50 ms by steps of 2 ms. For each pair of parameter values, the WC model was simulated at a sampling frequency of 10 kHz for 5 minutes. The resulting data were then downsampled to 1 kHz. As excitatory pyramidal cells contribute most strongly to EEG/MEG/ECoG signals, we associate activity in the excitatory populations of the model with signals in experimental data [81]. Simulated ICMs from the neuronal activity were then extracted in a similar way as for the empirical data from a time-frequency decomposition of the simulated signals.

### Spatial autoregressive model

The SAR model assumes that fluctuating neuronal signals, x = {*x_i_*}, are related through a model of structural equations, which relies on expressing each signal as a linear function of the others and weighted by a global coupling factor *k*, leading to

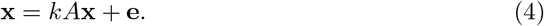

*A* represents the structural connectivity matrix, **e** is some additive noise that stands for the part of the signal that cannot be accounted for by SC. It is usually assumed to be normally distributed with zero mean and unknown covariance Σ, with spatial and temporal independence. We here further assumed that the variance of e is uniform across areas, reducing its covariance to a single constant value, Σ = *σ*^2^. According to Eq 4, x is multivariate normal with covariance

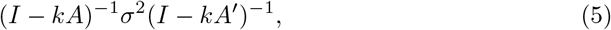

where *I* stands for the identity matrix, and’ is the regular matrix transposition. This model is also known as the simultaneous autoregressive model [36], which has been extensively used for the analysis of spatial data from diverse disciplines such as demography, economy and geography [82, 83].

We explored coupling parameter values from 0.001 to 0.999 by steps of 0.02. The SAR model provides a closed form for the covariance matrix that can be used to directly compute the predicted ICMs *via* its normalization. Of note, the SAR model does not provide frequency-resolved ICMs, we assume a common pattern across frequency bands.

### Model optimization

To evaluate the predictive power of the different models, we computed the Spearman correlation as well as distance between simulated and empirical ICM matrices. The distance (*D)* between two matrices *A* and *B* is defined as the root-mean-square deviation, also known as the Frobenius norm,

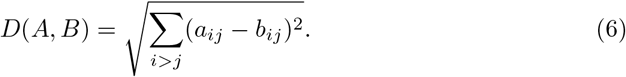

Optimal model parameters should maximize the similarity between simulated and empirical ICMs. This similarity was assessed using a simple objective function, representing the distance between simulations and empirical data:

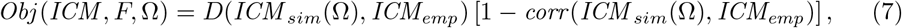

where *F* denoted the frequency band of interest, Ω the set of model’s parameters, *ICM_sim_*(Ω) the simulated ICM (either phase or envelope) according to the model’s parameters Ω, and *ICM_emp_* the corresponding empirical ICM measure. Optimal model parameters are those minimizing the sum of the objective functions averaged across all frequency bands:

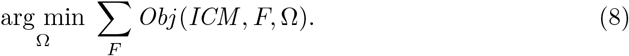

We explored different strategies of optimization by either minimizing the objective functions averaged over all ICM measures combining both original and lagged measures (one parameter set; *all*), over only original (*original*) or only lagged (*lagged*) ICM measures (two parameter sets), or for each ICM measure individually (four parameter sets; *individual*).

### Structural walks

Spectral graph theory relates the *k*-th power of the structural connectivity matrix A with walks of length *k*, such that the sum of the weights of all walks of length *k* corresponds to the entries of *A^k^*. In order to investigate whether topological information as reflected in the walks can predict empirical ICMs, we used a general linear model, a weighted sum of the powers of A, to fit phase and envelope ICMs separately and for each frequency band.

### Statistics

In order to probe the structure-function relationships, we computed the Spearman correlation between SC and phase and envelope ICM values for each frequency band, each animal and the group average. Additionally, we explored how much of the structure-function relationships may be explained by distance (fiber length). We computed the partial Spearman correlations between SC and phase and envelope ICM values, controlling for the distance. We also computed marginal Spearman correlations of both SC and ICMs with the distance. Similarity between phase and envelope ICMs was quantified by means of the Spearman correlation between the respective patterns for each frequency band, each animal and the group average. We computed also, for each type of ICM, the interfrequency consistency, i.e., the similarity between all pairs of frequency bands within each animal and the group average, and the interindividual consistency, i.e., the similarity between all pairs of animals within each frequency band. Since all brain connectivity measures are symmetrical, all correlation coefficients were computed using the upper triangular part of each connectivity matrix for the different scenarios.

In order to test the significance of the correlation coefficients and the significance of the difference between correlation coefficients (e.g., the difference between the correlation of pICMs with SC compared to the correlation of eICMs with SC) we used a frequentist approach based on bootstrap resampling [84]. For each correlation coefficient or pair of correlation coefficients, we generated 1 000 surrogate coefficients by randomly sampling connectivity values with replacement. Then, the overlap of the bootstrap distribution with zero was used to estimate the corresponding p-values. All statistical tests were rejected at *p* < 0.01 significance. Correction for multiple comparisons was performed by controlling the false discovery rate when appropriate [85].

## Acknowledgements

This work was supported by grants from the Deutsche Forschungsgemeinschaft (SFB936/A1/A2/Z3, SPP1665/EN533/13-1 and SPP2041/HI1286/6-1/EN533/15-1), the Human Brain Project (SGA2), and from the 2015 FLAG-ERA Joint Transnational Call for project FIIND (ANR-15-HBPR-0005). The funders had no role in study design, data collection and analysis, decision to publish, or preparation of the manuscript.

## Supporting information

**Fig S1.**
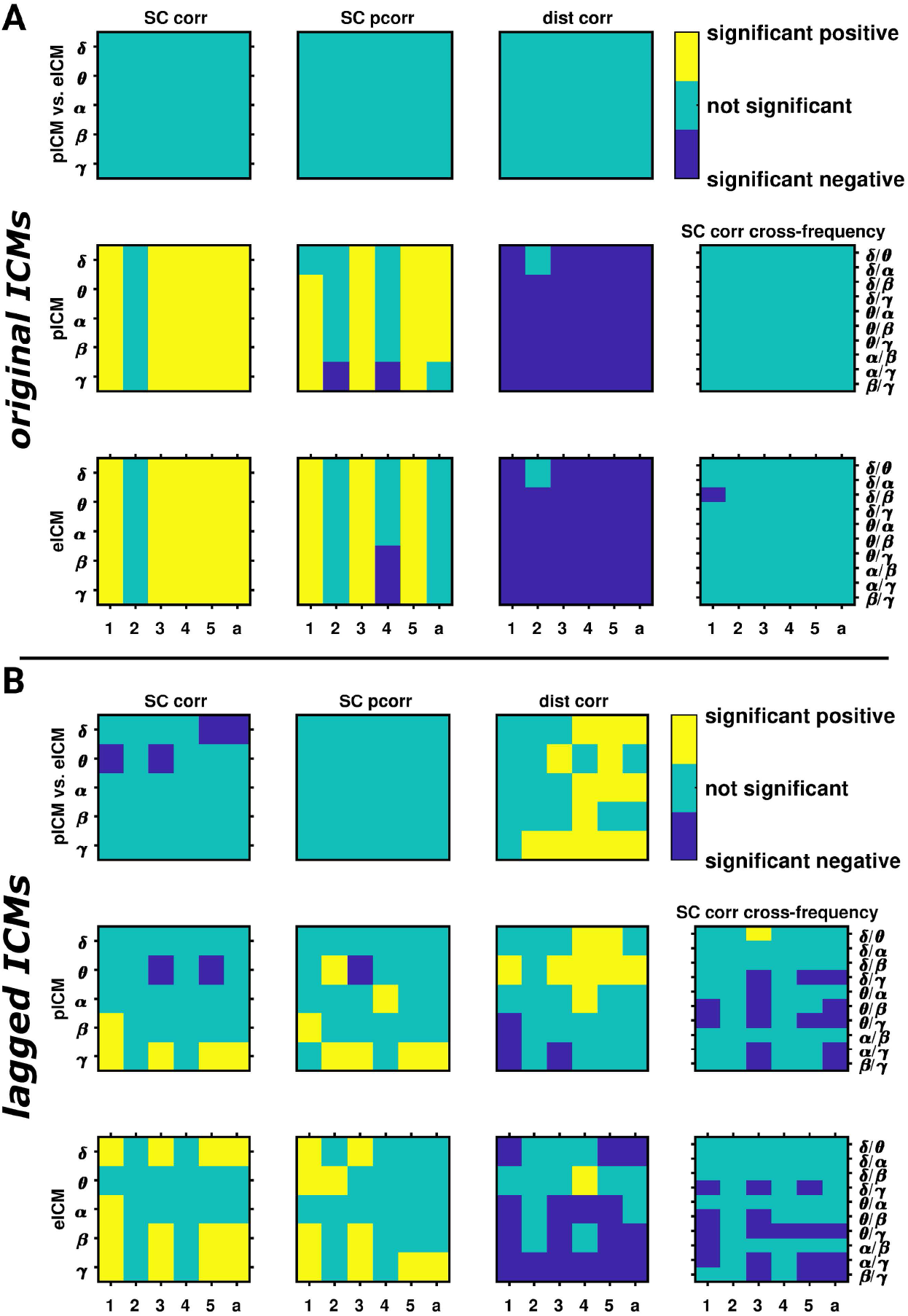
Structure-function relationships, statistics. (A) Significance of correlation values between structural connectivity and phase and envelope ICMs across frequency bands and animals. For each frequency band, each animal and the group average (a), we represented the significance of the difference in correlation (pICMs vs. eICMs) or the significance of the correlation (pICMs or eICMs) between SC and ICMs (SC corr), between SC and ICMs when controlling for the distance (SC pcorr), and between distance and ICMs (dist corr). We also represented the significance of the difference in correlation of SC with ICMs across frequency bands (SC corr cross-frequency). (B) Same as (A), when using lagged ICM measures.

**Fig S2.**
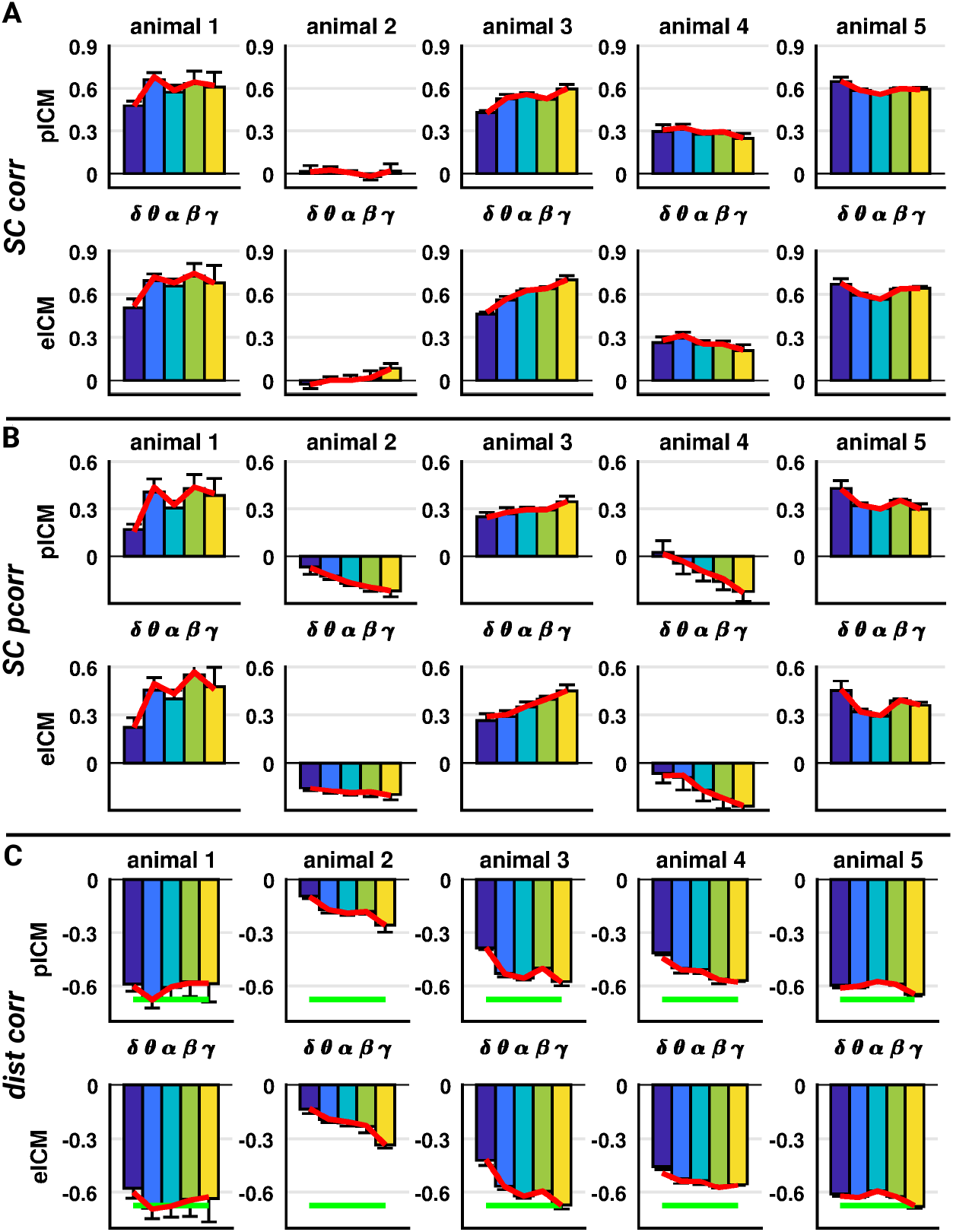
Structure-function relationships across animals when using original ICM measures. (A) Correlation between structural connectivity and phase and envelope ICMs across frequency bands, animals, and sessions when using original ICM measures. For each ICM measure, each animal and each frequency band, we represented the individual session correlations (bar chart representing means and associated standard deviations), as well as the correlation for the average session (red curve), between SC and ICMs (SC corr). (B) Same as (A) when controlling for the distance (SC pcorr). (C) Same as (A) when correlating ICMs with distance (dist corr). The green line represents the correlation between distance and SC.

**Fig S3.**
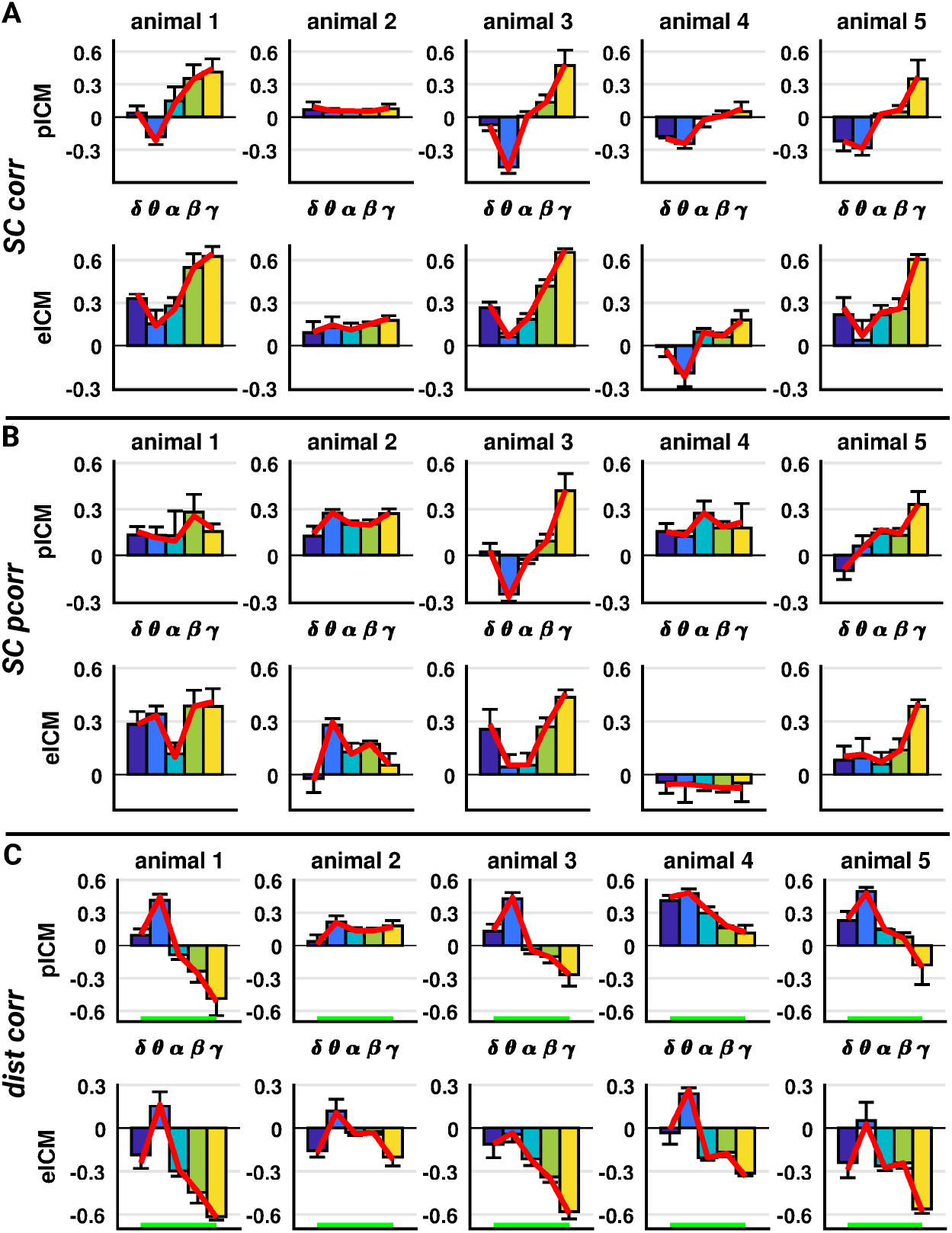
Structure-function relationships across animals when using lagged ICM measures. (A) Correlation between structural connectivity and phase and envelope ICMs across frequency bands, animals, and sessions when using lagged ICM measures. For each ICM measure, each animal and each frequency band, we represented the individual session correlations (bar chart representing means and associated standard deviations), as well as the correlation for the average session (red curve), between SC and ICMs (SC corr). (B) Same as (A) when controlling for the distance (SC pcorr). (C) Same as (A) when correlating ICMs with distance (dist corr). The green line represents the correlation between distance and SC.

**Fig S4.**
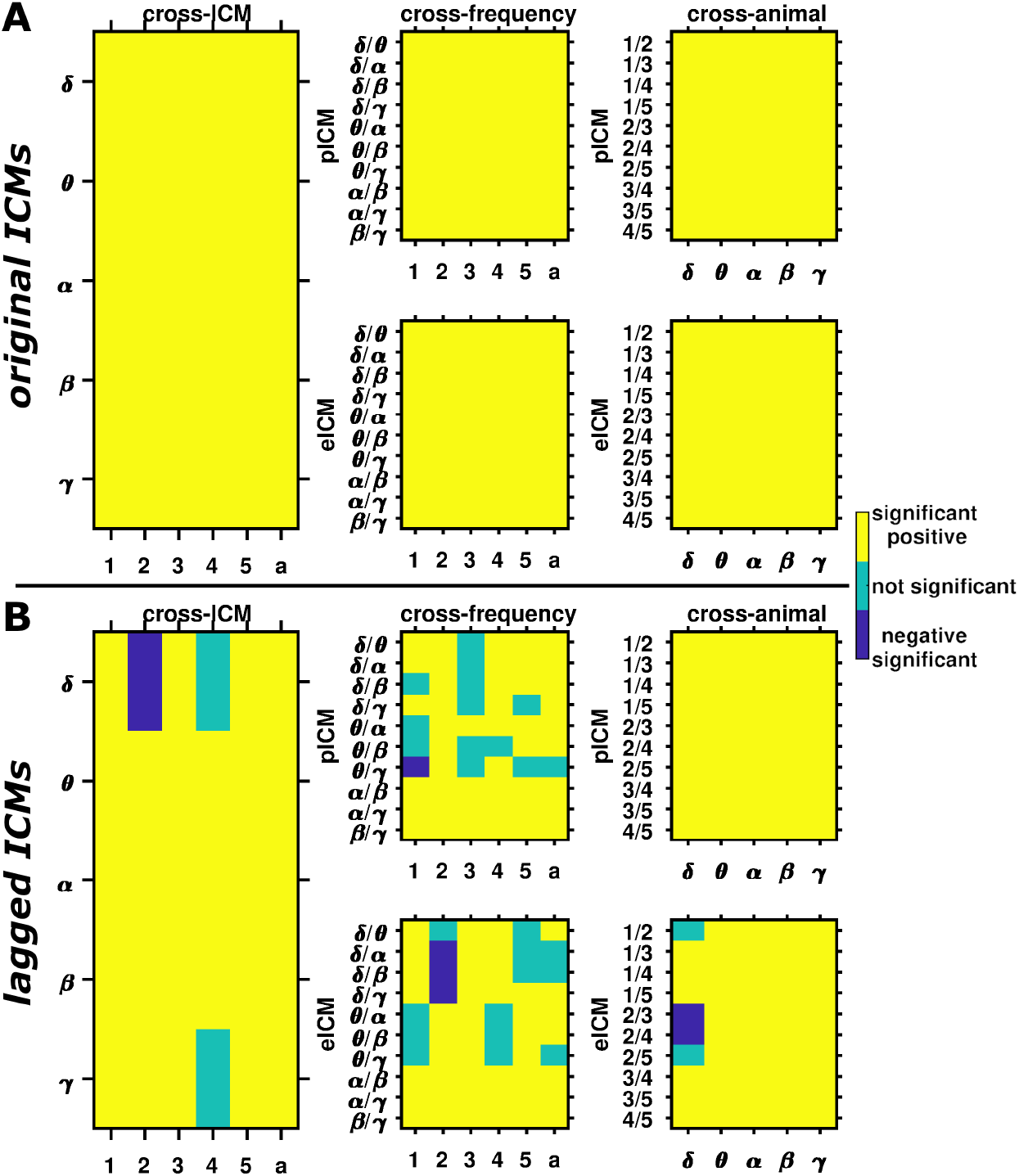
ICM similarity and consistency, statistics. (A) Significance of the similarity of phase and envelope ICMs, and interfrequency and interindividual consistency across frequency bands and animals. We represented the significance of the correlation between pICM and eICM (cross-ICM), for each frequency band, each animal and the group average (a); the significance of the correlation between ICMs across frequency bands (cross-frequency), for each ICM measure, each animal and the group average (a), and each pair of frequency bands; and the significance of the correlation between ICMs across animals (cross-animals), for each ICM measure, each frequency band and each pair of animals. (B) Same as (A) when using lagged ICM measures.

**Fig S5.**
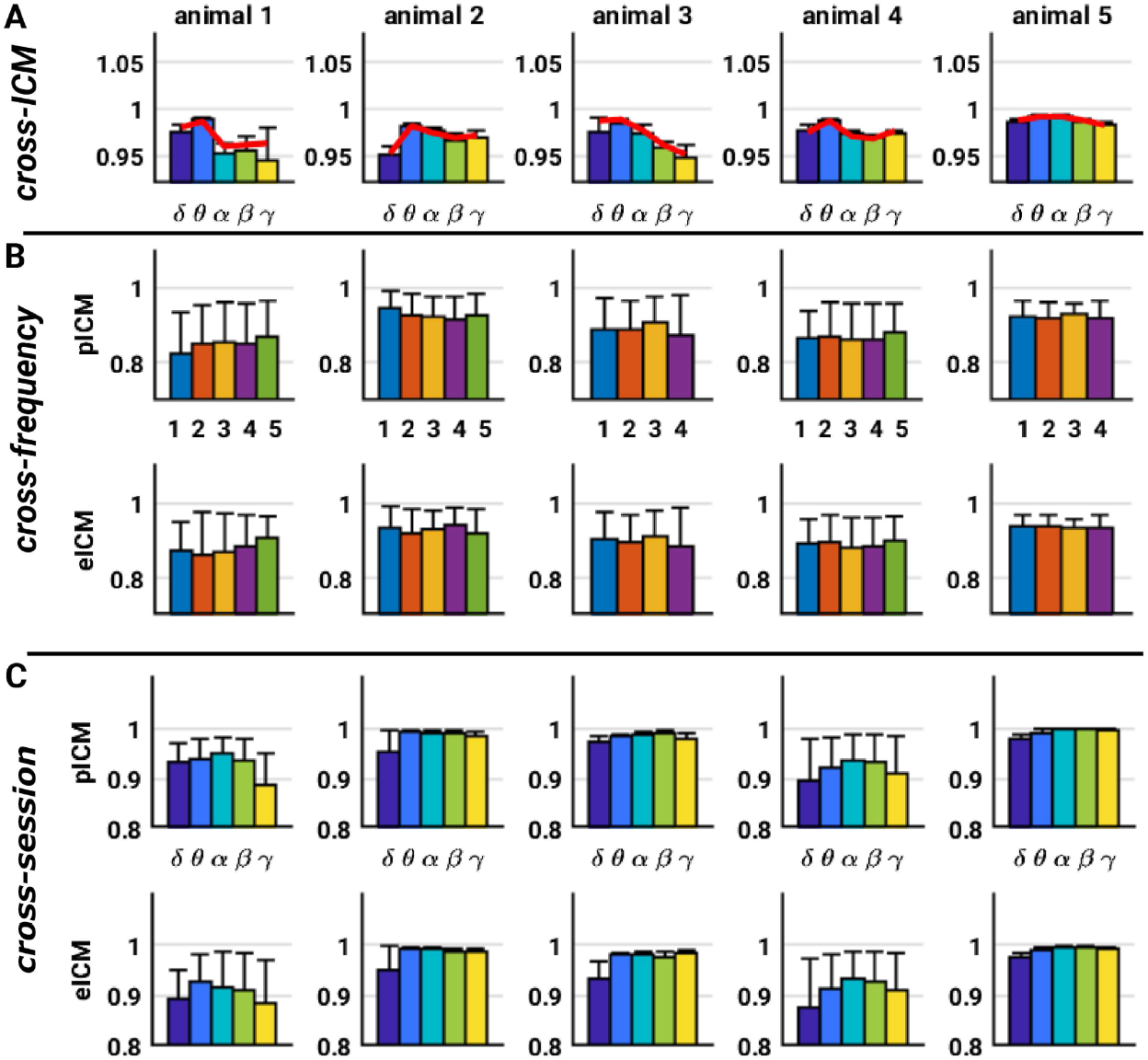
ICM similarity and consistency across animals when using original ICM measures. (A) Correlation between phase and envelope ICMs. For each animal and each frequency band, we represented the individual session correlations (bar chart representing means and associated standard deviations), as well as the correlation for the average session (red curve). (B) Interfrequency consistency of ICM patterns. For each animal, each ICM measure, and each session, we represented all pairwise correlations between frequency bands (bar chart representing means and associated standard deviations). (C) Intraindividual consistency of ICMs measures. For each animal, each ICM measure and each frequency band, we represented all pairwise correlations between sessions (bar chart representing means and associated standard deviations).

**Fig S6.**
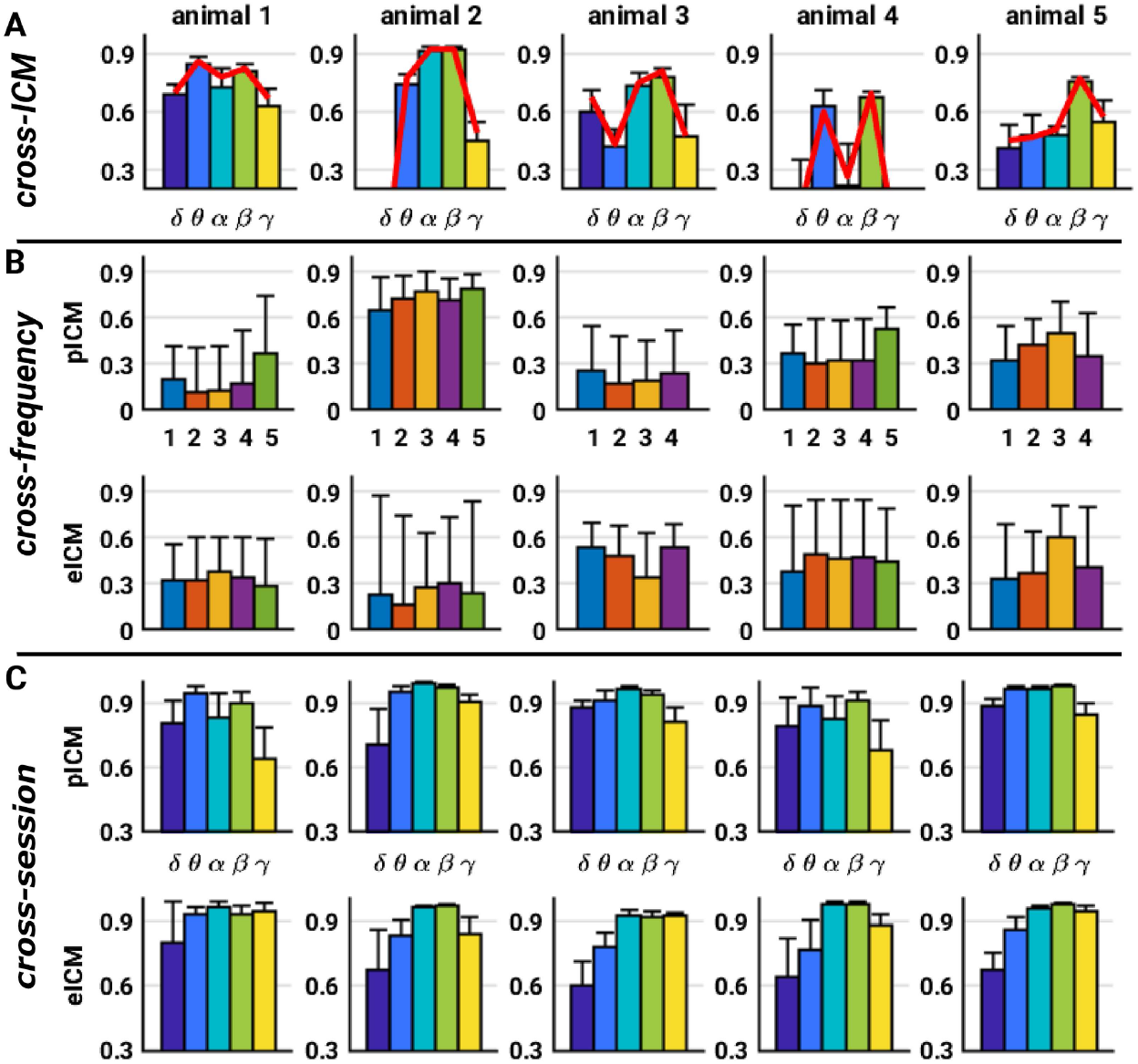
ICM similarity and consistency across animals when using lagged ICM measures. (A) Correlation between phase and envelope ICMs. For each animal and each frequency band, we represented the individual session correlations (bar chart representing means and associated standard deviations), as well as the correlation for the average session (red curve). (B) Interfrequency consistency of ICM patterns. For each animal, each ICM measure, and each session, we represented all pairwise correlations between frequency bands (bar chart representing means and associated standard deviations). (C) Intraindividual consistency of ICMs measures. For each animal, each ICM measure and each frequency band, we represented all pairwise correlations between sessions (bar chart representing means and associated standard deviations).

**Fig S7.**
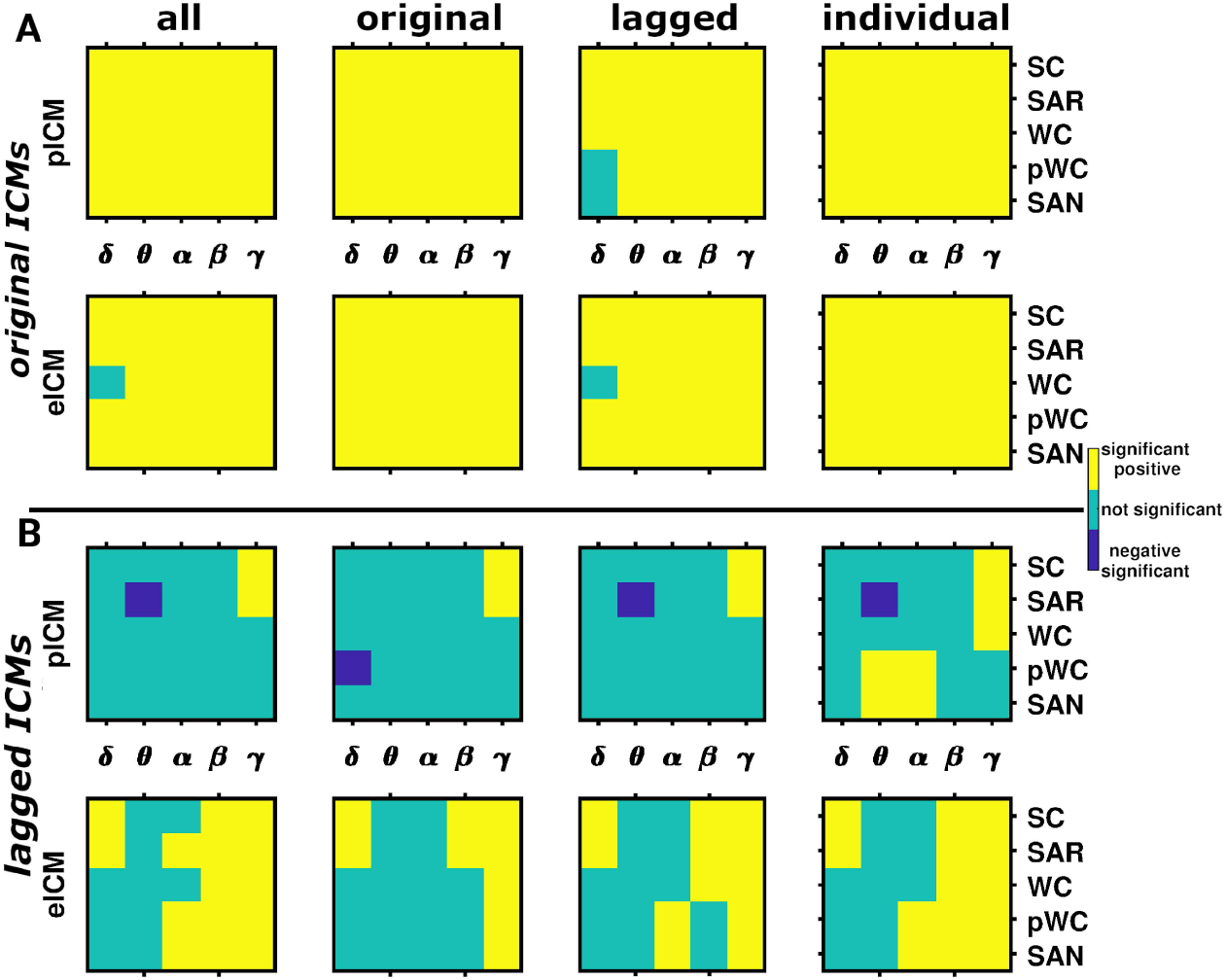
Predictive power of the computational models, statistics part 1. (A) Significance of the correlation between simulated and empirical phase and envelope ICMs across frequency bands. For each frequency band, each ICM measure, and each optimization strategy (columns), we represented the significance of the correlation between simulated and empirical ICM measures (including SC). (B) Same as (A) when using lagged ICM measures.

**Fig S8.**
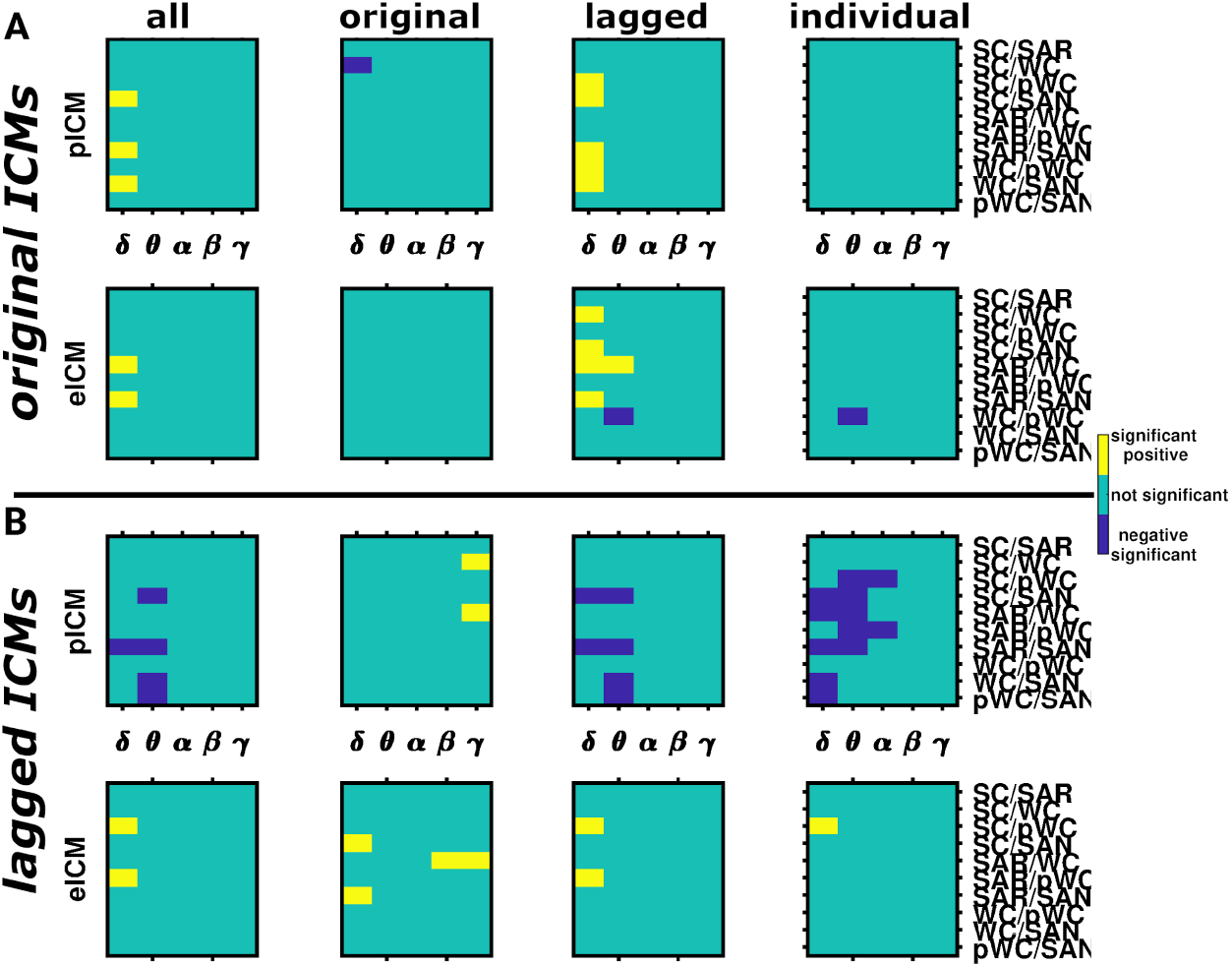
Predictive power of the computational models, statistics part 2. (A) Significance of the difference between models in the correlation between simulated and empirical phase and envelope ICMs across frequency bands. For each frequency band, each ICM measure, and each optimization strategy (columns), we represented the significance of the difference between all pairs of models (including SC) in the correlation between simulated and empirical ICM measures. (B) Same as (A) when using lagged ICM measures.

**Fig S9.**
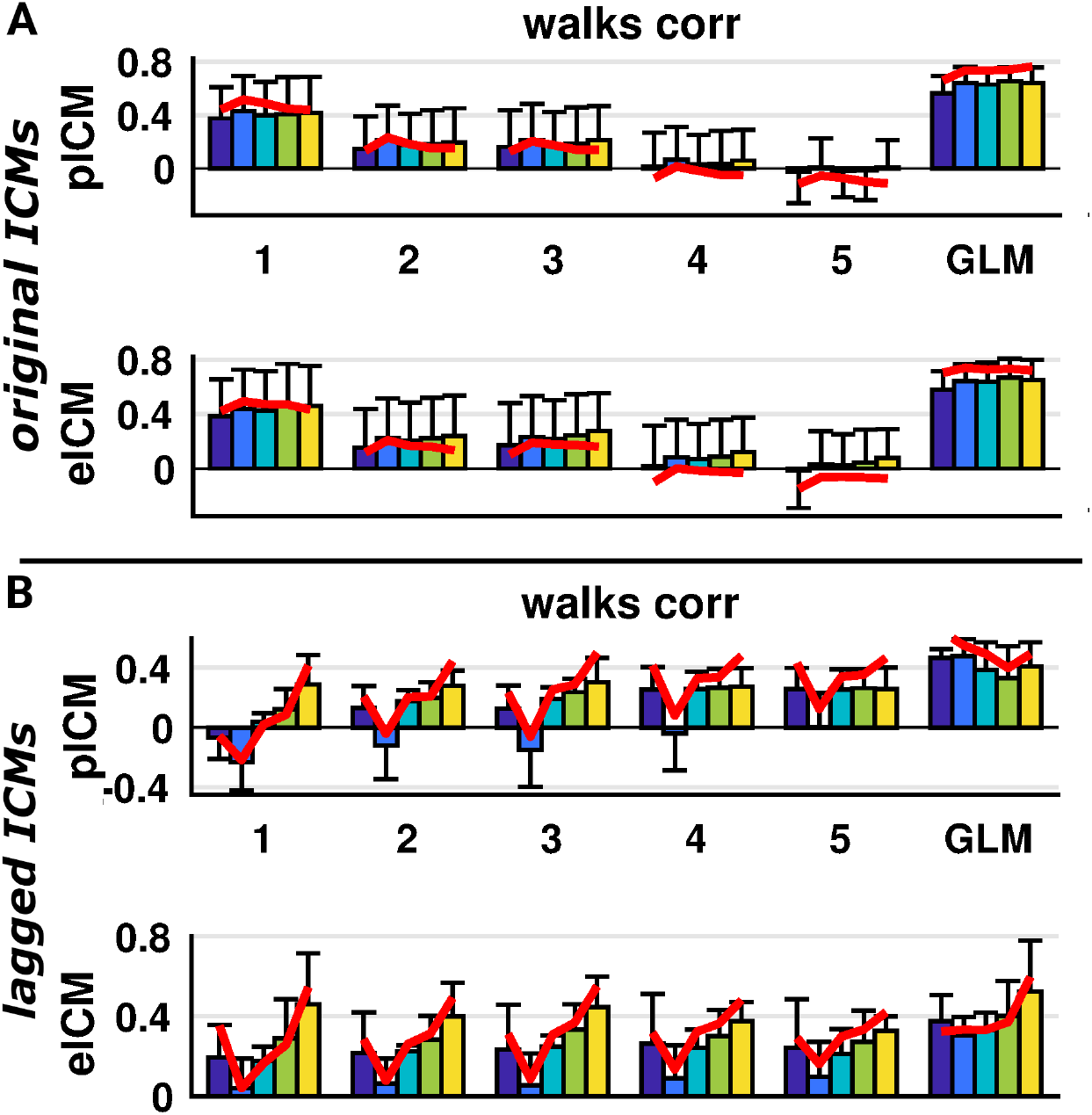
Predictive power of the structural walks. (A) Correlation between structural walks and empirical phase and envelope ICMs across frequency bands. For each walk length (and their linear combination) and each frequency band, we represented the individual correlations (bar chart representing means and associated standard deviations), as well as the correlation for the group average (red curve). (B) Same as (A) when using lagged ICM measures.

**Fig S10.**
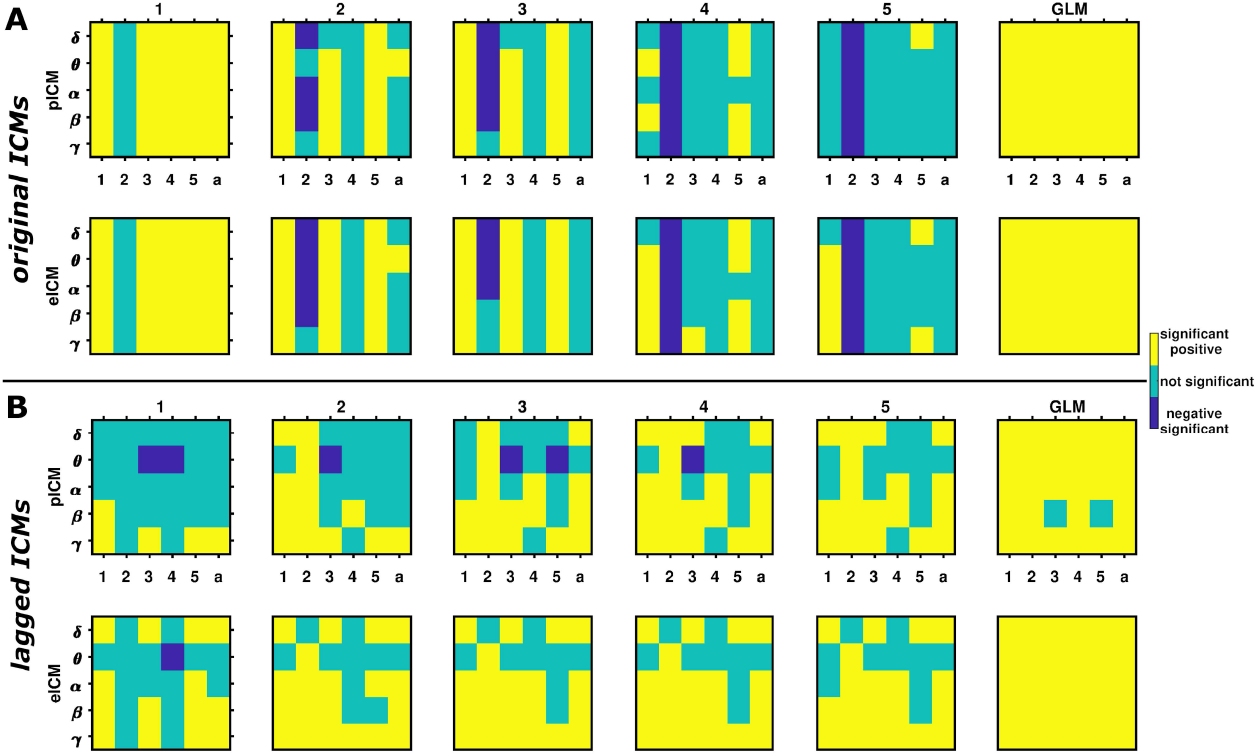
Predictive power of the structural walks, statistics. (A) Significance of the correlation between structural walks and empirical phase and envelope ICMs across frequency bands. For each frequency band, each ICM measure, and each walk length (and their combination), we represented the significance of the correlation between structural walks and empirical ICM measures. (B) Same as (A) when using lagged ICM measures.

## References

1. Engel AK, Gerloff C, Hilgetag CC, Nolte G. Intrinsic coupling modes: multiscale interactions in ongoing brain activity. Neuron. 2013;80(4):867–886. doi:10.1016/j.neuron.2013.09.038.

2. Fries P. A mechanism for cognitive dynamics: neuronal communication through neuronal coherence. Trends in Cognitive Sciences. 2005;9:474–480. doi:10.1016/j.tics.2005.08.011.

3. Fox M, Raichle ME. Spontaneous fluctuations in brain activity observed with functional magnetic resonance imaging. Nature Reviews Neuroscience. 2007;8:700–711. doi:10.1038/nrn2201.

4. Hipp JF, Engel AK, Siegel M. Oscillatory synchronization in large-scale cortical networks predicts perception. Neuron. 2011;69(2):387–396. doi:10.1016/j.neuron.2010.12.027.

5. Smith S, Nichols T, Vidaurre D, Winkler A, Behrens T, Glasser M, et al. A positive-negative mode of population covariation links brain connectivity, demographics and behavior. Nature Neuroscience. 2015;18(11):1565–1567. doi:10.1038/nn.4125.

6. Schnitzler A, Gross J. Normal and pathological oscillatory communication in the brain. Nature Reviews Neuroscience. 2005;6(4):285–296. doi:10.1038/nrn1650.

7. Uhlhaas P, Singer W. Neuronal dynamics and neuropsychiatric disorders: toward a translational paradigm for dysfunctional large-scale networks. Neuron. 2012;75(6):963–980. doi:10.1016/j.neuron.2012.09.004.

8. Fornito A, Bullmore ET. Connectomics: a new paradigm for understanding brain disease. European Neuropsychopharmacology. 2015;25(5):733–748. doi:10.1016/j.euroneuro.2014.02.011.

9. Honey CJ, Thivierge JP, Sporns O. Can structure predict function in the human brain? NeuroImage. 2010;52:766–776. doi:10.1016/j.neuroimage.2010.01.071.

10. Park HJ, Friston K. Structural and functional brain networks: from connections to cognition. Science. 2013;342(6158):1238411. doi:10.1126/science.1238411.

11. Deco G, Jirsa V, McIntosh A. Emerging concepts for the dynamical organization of resting-state activity in the brain. Nature Reviews Neuroscience. 2011;12:43–56. doi:10.1038/nrn2961.

12. Hipp JF, Hawellek DJ, Corbetta M, Siegel M, Engel AK. Large-scale cortical correlation structure of spontaneous oscillatory activity. Nature Neuroscience. 2012;15:884–890. doi:10.1038/nn.3101.

13. Siegel M, Donner T, Engel AK. Spectral fingerprints of large-scale neuronal interactions. Nature Reviews Neuroscience. 2012;13(2):121–134. doi:10.1038/nrn3137.

14. Singer W. Cortical dynamics revisited. Trends in Cognitive Sciences. 2013;17:616–626. doi:10.1016/j.tics.2013.09.006.

15. Sporns O. Contributions and challenges for network models in cognitive neuroscience. Nature Neuroscience. 2014;17(5):652–660. doi:10.1038/nn.3690.

16. Foster BL, He BJ, Honey CJ, Jerbi K, Maier A, Saalmann YB. Spontaneous Neural Dynamics and Multi-scale Network Organization. Frontiers in Systems Neuroscience. 2016;10:7. doi:10.3389/fnsys.2016.00007.

17. Le Bihan D, Heidi JB. Diffusion MRI at 25: exploring brain tissue structure and function. NeuroImage. 2011;61(2):324–341. doi:10.1016/j.neuroimage.2011.11.006.

18. Raichle M. Two views of brain function. Trends in Cognitive Sciences. 2010;14:180–190. doi:10.1016/j.tics.2010.01.008.

19. van den Heuvel MP, Hulshoff Pol HE. Exploring the brain network: A review on resting-state fMRI functional connectivity. European Neuropsychopharmacology. 2010;20:519–534. doi:10.1016/j.euroneuro.2010.03.008.

20. Koch MA, Norris DG, Hund-Georgiadis M. An investigation of functional and anatomical connectivity using magnetic resonance imaging. NeuroImage. 2002; 16(1):241–250. doi:10.1006/nimg.2001.1052.

21. Vincent JL, Patel GH, Fox MD, Snyder AZ, Baker JT, Van Essen DC, et al. Intrinsic functional architecture in the anaesthetized monkey brain. Nature. 2007;447:83–86. doi:10.1038/nature05758.

22. Hagmann P, Cammoun L, Gigandet X, Meuli R, Honey C, Wedeen V, et al. Mapping the structural core of human cerebral cortex. PLoS Biology. 2008;6:e159. doi:10.1371/journal.pbio.0060159.

23. Greicius MD, Supekar K, Menon V, Dougherty RF. Resting-state functional connectivity reflects structural connectivity in the default mode network. Cerebral Cortex. 2009;19:72–78. doi:10.1093/cercor/bhn059.

24. Chu C, Tanaka N, Diaz J, Edlow B, Wu O, Hämäläinen M, et al. EEG functional connectivity is partially predicted by underlying white matter connectivity. NeuroImage. 2015;108:23–33. doi:10.1016/j.neuroimage.2014.12.033.

25. Meier J, Tewarie P, Hillebrand A, Douw L, van Dijk BW, Stufflebeam SM, et al. A mapping between structural and functional brain networks. Brain Connectivity. 2016;6(4):298–311. doi:10.1089/brain.2015.0408.

26. Haüfe S, Nikulin VV, Muller KR, Nolte G. A critical assessment of connectivity measures for EEG data: A simulation study. NeuroImage. 2013;64:120–133. doi:10.1016/j.neuroimage.2012.09.036.

27. Palva JM, Palva S. Discovering oscillatory interaction networks with M/EEG: challenges and breakthroughs. Trends in Cognitive Sciences. 2012;16:219–230. doi:10.1016/j.tics.2012.02.004.

28. Finger H, Bönstrup M, Cheng B, Messé A, Hilgetag C, Thomalla G, et al. Modeling of large-scale functional brain networks based on structural connectivity from DTI: comparison with EEG derived phase coupling networks and evaluation of alternative methods along the modeling path. PLoS Computational Biology. 2016;12(8):e1005025. doi:10.1371/journal.pcbi.1005025.

29. Deco G, Jirsa V, Robinson PA, Breakspear M, Friston K. The dynamic brain: from spiking neurons to neural masses and cortical fields. PLoS Computational Biology. 2008;4:e1000092. doi:10.1371/journal.pcbi.1000092.

30. Deco G, Jirsa V, McIntosh AR, Sporns O, Kotter R. Key role of coupling, delay, and noise in resting brain fluctuations. Proceedings of the National Academy of Sciences of the USA. 2009;106:10302–10307.

31. Breakspear M. Dynamics models of large-scale brain activity. Nature Neuroscience. 2017;20:340–352. doi:10.1038/nn.4497.

32. Cabral J, Kringelbach ML, Deco G. Exploring the network dynamics underlying brain activity at rest. Progress in Neurobiology. 2014;114:102–131. doi:10.1016/j.pneurobio.2013.12.005.

33. Messíe A, Rudrauf D, Benali H, Marrelec G. Relating structure and function in the human brain: relative contributions of anatomy, stationary dynamics, and non-stationarities. PLoS Computational Biology. 2014;10:e1003530. doi:10.1371/journal.pcbi.1003530.

34. Messé A, Rudrauf D, Giron A, Marrelec G. Predicting functional connectivity from structural connectivity via computational models using MRI: an extensive comparison study. NeuroImage. 2015;111:65–75. doi:10.1016/j.neuroimage.2015.02.001.

35. Goñi J, van den Heuvel MP, Avena-Koenigsberger A, Velez de Mendizabal N, Betzel RF, Griffa A, et al. Resting-brain functional connectivity predicted by analytic measures of network communication. Proceedings of the National Academy of Sciences of the USA. 2014;111(2):833–838. doi:10.1073/pnas.1315529111.

36. Messé A, Benali H, Marrelec G. Relating structural and functional connectivity in MRI: a simple model for a complex brain. IEEE Transactions on Medical Imaging. 2015;34:27–37. doi:10.1109/TMI.2014.2341732.

37. Cabral J, Luckhoo H, Woolrich M, Joensson M, Mohseni H, Baker A, et al. Exploring mechanisms of spontaneous functional connectivity in MEG: how delayed network interactions lead to structured amplitude envelopes of band-pass filtered oscillations. NeuroImage. 2014;90:423–435.

38. Abeysuriya RG, Hadida J, Sotiropoulos SN, Jbabdi S, Becker R, Hunt BA, et al. A biophysical model of dynamic balancing of excitation and inhibition in fast oscillatory large-scale networks. PLoS Computational Biology. 2018;14:1–27. doi:10.1371/journal.pcbi.1006007.

39. Tewarie P, Abeysuriya R, Byrne A, O’Neill GC, Sotiropoulos SN, Brookes MJ, et al. How do spatially distinct frequency specific MEG networks emerge from one underlying structural connectome? The role of the structural eigenmodes. NeuroImage. 2019;186:211–220. doi:10.1016/j.neuroimage.2018.10.079.

40. Stitt I, Hollensteiner KJ, Galindo-Leon E, Pieper F, Fiedler E, Stieglitz T, et al. Dynamic reconfiguration of cortical functional connectivity across brain states. Scientific Reports. 2017;7:8797. doi:10.1038/s41598-017-08050-6.

41. Delettre C, Mess’ A, Dell LA, Foubet O, Heuer K, Larrat B, et al. Comparison between diffusion MRI tractography and histological tract-tracing of cortico-cortical structural connectivity in the ferret brain. Network Neuroscience. 2019;3:1038–1050. doi:10.1162/netn_a_00098.

42. Bizley JK, King AJ. Visual influences on ferret auditory cortex. Hearing Research. 2009;258:55–63. doi:https://doi.org/10.1016/j.heares.2009.06.017.

43. Messé A. Parcellation influence on the connectivity-based structure-function relationship in the human brain. Human Brain Mapping. in press;doi:10.1002/hbm.24866.

44. Straathof M, Sinke MR, Dijkhuizen RM, Otte WM. A systematic review on the quantitative relationship between structural and functional network connectivity strength in mammalian brains. Journal of Cerebral Blood Flow and Metabolism. 2019;39:189–209. doi:10.1177/0271678X18809547.

45. Wirsich J, Ridley B, Bésson P, Jirsa V, Benar C, Ranjeva JP, et al. Complementary contributions of concurrent EEG and fMRI connectivity for predicting structural connectivity. NeuroImage. 2017;161:251–260. doi:10.1016/j.neuroimage.2017.08.055.

46. Tewarie P, Hillebrand A, van Dellen E, Schoonheim M, Barkhof F, Polman C, et al. Structural degree predicts functional network connectivity: a multimodal resting-state fMRI and MEG study. NeuroImage. 2014;97:296–307. doi:10.1016/j.neuroimage.2014.04.038.

47. Garcés P, Pereda E, Hernàndez-Tamames JA, Del-Pozo F, Maestù F, Angel Pineda-Pardo J. Multimodal description of whole brain connectivity: a comparison of resting state MEG, fMRI, and DWI. Human Brain Mapping. 2016;37(1):20–34. doi:10.1002/hbm.22995.

48. Colclough GL, Woolrich MW, Tewarie PK, Brookes MJ, Quinn AJ, Smith SM. How reliable are MEG resting-state connectivity metrics? NeuroImage. 2016;138:284 – 293. doi:https://doi.org/10.1016/j.neuroimage.2016.05.070.

49. Bastos AM, Schoffelen JM. A Tutorial Review of Functional Connectivity Analysis Methods and Their Interpretational Pitfalls. Frontiers in Systems Neuroscience. 2016;9:175. doi:10.3389/fnsys.2015.00175.

50. Palva JM, Wang SH, Palva S, Zhigalov A, Monto S, Brookes MJ, et al. Ghost interactions in MEG/EEG source space: A note of caution on inter-areal coupling measures. NeuroImage. 2018;173:632–643. doi:10.1016/j.neuroimage.2018.02.032.

51. Siems M, Siegel M. Dissociated neuronal phase- and amplitude-coupling patterns in the human brain. NeuroImage. 2020;209:116538. doi:10.1016/j.neuroimage.2020.116538.

52. Galindo-Leon EE, Stitt I, Pieper F, Stieglitz T, Engler G, Engel AK. Context-specific modulation of intrinsic coupling modes shapes multisensory processing. Science Advances. 2019;5(4). doi:10.1126/sciadv.aar7633.

53. Messé A, Hütt MT, Hilgetag CC. Toward a theory of coactivation patterns in excitable neural networks. PLoS Computational Biology. 2018;14:1–19. doi:10.1371/journal.pcbi.1006084.

54. Cabral J, Kringelbach ML, Deco G. Functional connectivity dynamically evolves on multiple time-scales over a static structural connectome: Models and mechanisms. NeuroImage. 2017;160:84–96. doi:10.1016/j.neuroimage.2017.03.045.

55. Honey CJ, Sporns O, Cammoun L, Gigandet X, Thiran JP, Meuli R, et al. Predicting human resting-state functional connectivity from structural connectivity. Proceedings of the National Academy of Sciences of the USA. 2009;106:2035–2040. doi:10.1073/pnas.0811168106.

56. Marrelec G, Messé A, Giron A, Rudrauf D. Functional connectivity’s degenerate view of brain computation. PLoS Computational Biology. 2016;12(10):e1005031. doi:10.1371/journal.pcbi.1005031.

57. Nolte G, Bai O, Wheaton L, Mari Z, Vorbach S, Hallett M. Identifying true brain interaction from EEG data using the imaginary part of coherency. Clinical Neurophysiology. 2004;115(10):2292–2307. doi:10.1016/j.clinph.2004.04.029.

58. Brookes MJ, Woolrich MW, Barnes GR. Measuring functional connectivity in MEG: A multivariate approach insensitive to linear source leakage. NeuroImage. 2012;63:910 – 920. doi:10.1016/j.neuroimage.2012.03.048.

59. Engel AK, Fries P, Singer W. Dynamic predictions: oscillations and synchrony in top-down processing. Nature Reviews Neuroscience. 2001;2(10):704–716. doi:10.1038/35094565.

60. Engel A, Konig P, Kreiter A, Singer W. Interhemispheric synchronization of oscillatory neuronal responses in cat visual cortex. Science. 1991;252(5009):1177–1179. doi:10.1126/science.252.5009.1177.

61. Sjøgård M, Bourguignon M, Costers L, Dumitrescu A, Coolen T, Roshchupkina L, et al. Intrinsic/extrinsic duality of large-scale neural functional integration in the human brain. bioRxiv. 2020;doi:10.1101/2020.04.21.053579.

62. Deco G, Cruzat J, Cabral J, Knudsen GM, Carhart-Harris RL, Whybrow PC, et al. Whole-brain multimodal neuroimaging model using serotonin receptor maps explains non-linear functional effects of LSD. Current Biology. 2018;28:3065–3074. doi:10.1016/j.cub.2018.07.083.

63. Schmidt M, Bakker R, Shen K, Bezgin G, Diesmann M, van Albada SJ. A multi-scale layer-resolved spiking network model of resting-state dynamics in macaque visual cortical areas. PLoS Computational Biology. 2018;14(10):1–38. doi:10.1371/journal.pcbi.1006359.

64. Rubehn B, Bosman C, Oostenveld R, Fries P, Stieglitz T. A MEMS-based flexible multichannel ECoG-electrode array. Journal of Neural Engineering. 2009;6(3):036003. doi:10.1088/1741-2560/6/3/036003.

65. Oostenveld R, Fries P, Maris E, Schoffelen JM. FieldTrip: open source software for advanced analysis of MEG, EEG, and invasive electrophysiological data. Computational Intelligence and Neuroscience. 2011;2011:156869. doi:10.1155/2011/156869.

66. Brookes MJ, Woolrich M, Luckhoo H, Price D, Hale JR, Stephenson MC, et al. Investigating the electrophysiological basis of resting state networks using magnetoencephalography. Proceedings of the National Academy of Sciences of the USA. 2011;108:16783–16788. doi:10.1073/pnas.1112685108.

67. de Pasquale F, Della Penna S, Snyder A, Marzetti L, Pizzella V, Romani G, et al. A cortical core for dynamic integration of functional networks in the resting human brain. Neuron. 2012;74(4):753–764. doi:10.1016/j.neuron.2012.03.031.

68. Veraart J, Novikov DS, Christiaens D, Ades-aron B, Sijbers J, Fieremans E. Denoising of diffusion MRI using random matrix theory. NeuroImage. 2016;142:394–406. doi:10.1016/j.neuroimage.2016.08.016.

69. Kellner E, Dhital B, Kiselev VG, Reisert M. Gibbs-ringing artifact removal based on local subvoxel-shifts. Magnetic Resonance in Medicine. 2016;76(5):1574–1581. doi:10.1002/mrm.26054.

70. Jenkinson M, Beckmann CF, Behrens TEJ, Woolrich MW, Smith SM. FSL. NeuroImage. 2012;62:782–790. doi:10.1016/j.neuroimage.2011.09.015.

71. Andersson JL, Sotiropoulos SN. An integrated approach to correction for off-resonance effects and subject movement in diffusion MR imaging. NeuroImage. 2016;125:1063–1078. doi:10.1016/j.neuroimage.2015.10.019.

72. Tustison NJ, Avants BB, Cook PA, Zheng Y, Egan A, Yushkevich PA, et al. N4ITK: improved N3 bias correction. IEEE Transactions on Medical Imaging. 2010;29(6):1310–1320. doi:10.1109/TMI.2010.2046908.

73. Jenkinson M, Bannister P, Brady M, Smith S. Improved optimization for the robust and accurate linear registration and motion correction of brain images. NeuroImage. 2002;17(2):825–841.

74. Christiaens D, Reisert M, Dhollander T, Sunaert S, Suetens P, Maes F. Global tractography of multi-shell diffusion-weighted imaging data using a multi-tissue model. NeuroImage. 2015;123:89–101. doi:10.1016/j.neuroimage.2015.08.008.

75. Jeurissen B, Tournier JD, Dhollander T, Connelly A, Sijbers J. Multi-tissue constrained spherical deconvolution for improved analysis of multi-shell diffusion MRI data. NeuroImage. 2014;103:411–426. doi:10.1016/j.neuroimage.2014.07.061.

76. Tournier JD, Calamante F, Connelly A. MRtrix: diffusion tractography in crossing fiber regions. International Journal of Imaging Systems and Technology. 2012;22(1):53–66. doi:10.1002/ima.22005.

77. Deco G, Jirsa VK. Ongoing cortical activity at rest: criticality, multistability, and ghost attractors. The Journal of Neuroscience. 2012;32:3366–3375. doi:10.1523/JNEUROSCI.2523-11.2012.

78. Wilson HR, Cowan JD. Excitatory and inhibitory interactions in localised populations of model neurons. Biophysical Journal. 1972;12:1–24.

79. Tononi G, Sporns O, Edelman G. A measure for brain complexity: relating functional segregation and integration in the nervous system. Proceedings of the National Academy of Sciences of the USA. 1994;91:5033–5037.

80. Cabral J, Hugues E, Sporns O, Deco G. Role of local network oscillations in resting-state functional connectivity. NeuroImage. 2011;57:130–139. doi:10.1016/j.neuroimage.2011.04.010.

81. Buzsáki G, Anastassiou CA, Koch C. The origin of extracellular fields and currents - EEG, ECoG, LFP and spikes. Nature Reviews Neuroscience. 2012;13:407–420. doi:10.1038/nrn3241.

82. Lesage JP, Olivier P. Bayesian model averaging for spatial econometric models. Geographical Analysis. 2007;39:241–267. doi:10.1111/j.1538-4632.2007.00703.x.

83. de Oliveira V, Song JJ. Bayesian analysis of simultaneous autoregressive models. Sankhya: The Indian Journal of Statistics. 2008;70:323–350.

84. Wilcox RR. Comparing dependent robust correlations. British Journal of Mathematical and Statistical Psychology. 2016;69(3):215–224. doi:10.1111/bmsp.12069.

85. Benjamini Y, Krieger AM, Yekutieli D. Adaptive linear step-up procedures that control the false discovery rate. Biometrika. 2006;93(3):491–507. doi:10.1093/biomet/93.3.491.

